# Asymmetrical cleavages of *Sleeping Beauty* transposons generate multiple excised transposon fragments during transposition

**DOI:** 10.1101/659086

**Authors:** Qilin Gu, Xiaojie Yang, Qing Li, Yong Long, Guili Song, Desheng Pei, Perry B. Hackett, Jun Chen, Jinrong Peng, Zongbin Cui

## Abstract

Although the *Sleeping Beauty* (SB) transposon is the most validated DNA transposon used as a gene delivery vehicle in vertebrates, many details of the excision and integration steps in the transposition process are unclear. We have probed in detail the products of the excision step and apparent selective integration of a subset of those products during transposition. The standard model of SB transposase-mediated transposition includes symmetrical cleavages at both ends of the transposon for excision and re-integration in another DNA sequence. In our analysis of excised transposon fragments (ETFs), we found evidence for the requirement of certain flanking sequences for efficient cleavage and a significant rate of asymmetrical cleavage during the excision process that generates multiple ETFs. Our results suggest that the cleavage step by SB transposase is not as precise as indicated in most models. Repair of the donor ends can produce eight footprint sequences (TACTGTA, TACAGTA, TACATA, TACGTA, TATGTA, TACTA, TAGTA and TATA). Our data also suggest that mismatch repair (MMR) is not an essential requirement for footprint formation. Among the twenty liberated ETFs, only eight appear to effectively re-integrate into TA sites distributed across the genome, supporting earlier findings of unequal rates of excision and reintegration during SB transposition. These findings may be important in considerations of efficiency of SB transposon remobilization, selection of TA integration sites and detection of SB excision and integration loci, all of which may be important in human gene therapy.

## INTRODUCTION

Members of Tc1/*mariner* superfamily can be found in various species from fungi to animals. Although they are among the most widespread transposons in vertebrates, nearly all are inactive due to mutations in the genes encoding their respective transposases (1). More than 20 years ago, a defective Tc1/*mariner* element found in salmon was resurrected to activity and named *Sleeping Beauty* (SB) (2). Several other restored transposons followed, including *Frog Prince* (3) and *Hsmar1* (4). Since then, the SB transposon system has been widely utilized for genome manipulations in vertebrates (5). Over the past two decades, the efficiency of the SB system has been improved by optimization of the transposon structure (6–8), including *T_2_*(7) used in this study and *T_4_* (8), and by amino acid substitutions in the original SB10 transposase (6,9–12) including SB11 that is featured in our studies and which is undergoing clinical trials in humans (13, 14).

The SB transposon, like other Tc1/*mariner* elements, transposes through a cut-and-paste mechanism by a DDE-type transposase (1). The transposable element is excised from its original location, the *donor* sequence, and generally re-integrates into a TA target sequence (1). The excision of a transposon from a donor DNA sequence begins with a single-strand nick to generate a free 3’-OH group followed by the cleavage of the complementary DNA strand. Most DDE-transposases use a single, active site to cleave both DNA strands to create a DNA-hairpin structure at the ends of the transposon (5). However, excision of Tc1*/mariner* elements, including Tc1 (15), *Himar1* (16) and *Mos1* (17, 18), involves a pair of staggered, double-strand DNA breaks (DSBs) to generate extrachromosomal, excised transposon fragments (ETFs). The staggered cuts of Tc1/*mariner* transposons generate overhangs that can form either 2-bp (Tc1, Tc3, *mariner*, and IS630 transposases (15, 19)) or 3-bp (SB (20)) footprints at the original donor sites. Recently, SB transposon inversion-circles that are products of autointegration have been detected, indicating that SB transposons can be fully excised prior to their integration into new sites (21).

The recognition and cleavage steps of Tc1/*mariner* transposases have been studied by dissecting the crystal structures of *Mos1* paired-end complexes (22–24). *Hsmar1* transposition is carried out by two sequential strand cleavages and one strand transfer reaction at the same transposon end (25). Tc1 and Tc3 transposases additionally recognize some bases adjacent to the target, which can influence the frequency of transposition into a particular TA site (26, 27). DNA structure at the insertion site can also influence the mobility of *mariner* (28) and SB elements (29–32). Although we assume that DDE-type transposases of the Tc1/*m*ariner superfamily recognize and cleave transposon ends in a similar manner, the molecular aspects underlying the recognition, cleavage and reintegration by SB transposase are poorly understood because i) *in vitro* SB transposition assays remain unsuccessful; ii) the crystal structure of full-length SB transposase has not been resolved, although the NMR structure of the SB11 DNA-binding domain (33) and crystal structure of the SB100X catalytic domain (11) have been reported; and iii) compared to other Tc1/*mariner* transposons, the inverted terminal repeat (ITR) sequences that comprise the termini of SB transposons are considerably longer and each has two transposase-binding, direct repeat (DR) sequences (Figure 1A) (7). Hence, the ITRs of SB transposons are called IR/DRs for their unusual structure. This last feature introduces several questions about the coordination of transposase enzymes in both the cleavage and integration reactions; e.g., whether dimers or tetramers form to affect either or both steps.

**Figure 1.**
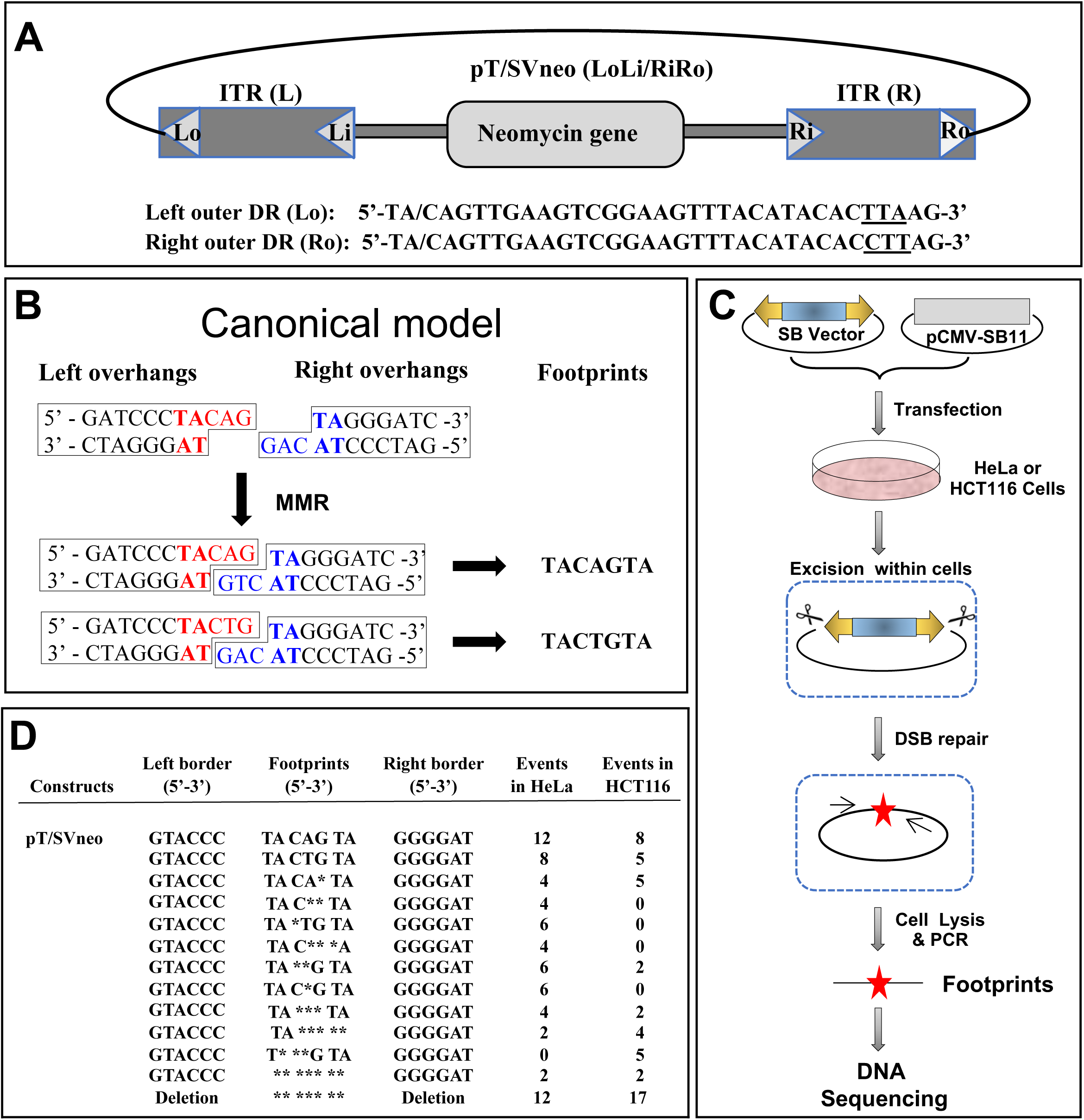
SB-mediated transposition and the products of transposon. (**A**) Structure of pT/SVneo with its classical inverted terminal repeats (ITRs) abbreviated as LoLi/RiRo. The *neomycin* gene expression cassette (neo) is flanked by the left and right ITRs that contain direct repeats (DRs). The sequences of the left and right outer DRs (Lo and Ro) are shown and the terminal sequences are complimentary (inverted). Flanking TA-dinucleotides are separated by a slash from the transposon outer DRs. The three different nucleotides in Lo and Ro are underlined. (**B**) The canonical SB-mediated excision model. In this model, the canonical 7-bp footprints result from the repair of broken donor ends by DNA mismatch repair (MMR). (**C**) Experimental design for footprint detection. (**D**). Footprints of excised pT/SVneo transposons. * indicates a missing base from canonical footprints.

In the standard model, SB transposon excision results from staggered cuts at both ends of the transposon that leaves a 3-bp single stranded overhanging sequence in both the donor DNA and the excised transposon. When the wounded DNA is repaired, either of two 7-bp canonical footprints, TAC<u>A</u>GTA and TAC<u>T</u>GTA, that differ in the fourth position (underlined) are left due to ambiguous repair of the mismatch at this position (Figure 1B) (20). The model suggests that SB transposase precisely cleaves the donor DNA and the targeted integration site to allow seamless integration of an excised transposon. However, if excision is not precise, i.e., the cleavage sites leave 3’-overhangs that are not exactly 3-bp, then such liberated transposons may not be candidates for re-entry into target DNA sites that were precisely cut. In order to further clarify the cleavage and re-integration steps by SB transposase, we constructed a large number of transposons that had varying terminal sequences (Figure S1). We then characterized in detail ETFs that appear to be liberated during excision and their respective re-integration abilities to complete the transposition process. These results refine the standard model of SB transposition and should be helpful for further understanding the transposition dynamics of SB and other Tc1/*mariner* members.

## MATERIAL AND METHODS

### Cell culture

HeLa and HCT116 cells were cultured in Dulbecco’s modified Eagle’s medium (DMEM) supplemented with 10% fetal bovine serum (FBS). XRS5 cells were cultured in Alpha minimum essential medium without ribonucleosides and deoxyribonucleosides with 2 mM L-glutamine supplemented with 10% FBS. M059J cells were grown in a medium containing a 1:1 mixture of DMEM and Ham’s F12 medium supplemented with 10% FBS.

### Zebrafish husbandry

Zebrafish (*Danio rerio*) were raised and maintained according to standard laboratory conditions. Our animal research protocol was approved by the Institutional Animal Care and Use Committee of the Institute of Hydrobiology.

### Plasmid constructs

The construct pT_2_/tiHsp70-SB11-SV40-Neo has a SB transposase gene that is driven by the tilapia heat-shock promoter 70 (tiHsp70) (34), and the G418-resistance gene (*neo*) that is controlled by the SV40 promoter. The plasmids we employed are named on the basis of their IR/DR sequences; Lo and Ro are the left and right outer DRs that are juxtaposed to the integration site whereas Li and Ri are the inner DRs that comprise the internal borders of the ITRs (Figure 1A). Plasmids LoLi/RiRo (pT/SVneo), pCMV-SB11, TAN_4_Lo, TAN_4_Ro, TAN_4_Lo/TAN_4_Ro, Lo(S3), LoLi/LiLo and RoRi/RiRo have been described previously (7). Ro(S3), TAN_4_Lo(S3), TAN_4_Ro(S3), TAN_4_Lo/Ro(S3), Lo(S3)/TAN_4_Ro and Lo(S3)/Ro(S3) were constructed with elements in LoLi/LiLo, RoRi/RiRo, pT/HindIIIneo, TAN_4_Lo, TAN_4_RoRi/RiRo, TAN_4_Lo, TAN_4_Ro(S3) and Lo(S3) (Figure S1). The primers used to generate these constructs are listed in Table S1.

The ETF capture constructs were reconstructed from the vector pT_2_/HB. Primer pairs used to generate these vectors are listed in Table S1 and the details are described in Tables S2 and S3.

The Blasticidin-resistant gene, *bsd,* in the vector pT_2_/zbHSP70-Bsd-SV40 was driven by the zebrafish HSP70 promoter (35). Twenty-eight vectors were designed to generate artificial ETFs by restriction endonuclease reactions (Table S4). The primers used to generate these vectors were listed in Table S1 and the details were described in Table S5. All vectors were confirmed by sequencing.

### Footprint detection assay

The pCMV-SB11 (100 ng) and pT/SVneo transposon (500 ng) plasmids were co-transfected into either HeLa or HCT116 cells (3×10^5^) in 35-mm culture dishes and incubated at 37°C for 3 days. Total DNA was extracted for footprint analysis as described previously (36). Primer pairs (1-For/1-Rev and 2-For/2-Rev in Table S1) were used for the first and second round PCR, respectively.

### Plasmid-rescue assays

The plasmid rescue assay was performed as previously described with minor modifications (21). The plasmid pT_2_/tiHsp70-SB11-SV40-rpsL, in which the SB11 transposase can be induced at 37°C under the control of tiHsp70 promoter, and expression of the streptomycin-sensitive gene *rpsL*, which is not streptomycin (Strep)-resistant (37), was controlled by the SV40 promoter. The plasmid contains the ampicillin (*amp*) gene. HeLa (MMR^+^) or HCT116 (MMR^-^) cells were transfected with pT_2_/tiHsp70-SB11-SV40-rpsL plasmids and cultured at 32 °C for 24 hours. Then, cells were either heat-shocked at 37°C for 2 hours or maintained at 32°C (no heat-shock). Plasmids were recovered from these cells three days post-transfection and subjected to T7 Endonuclease I (T7E) digestion. Digested DNA products were transformed into Top10 chemically competent *E. coli.* cells and subjected to a selection of either Amp (100 μg/ml) or a double selection of Amp/Strep (100 μg/ml and 30 μg/ml, respectively). The excision of transposon in plasmid pT_2_/tiHsp70-SB11-SV40-rpsL and DNA repair at the excision sites led to the generation of circularized plasmids without the *rpsL* gene in transfected HeLa and HCT116 cells. Amp^R^/Strep^R^-bacteria colonies were considered as “footprint colonies” and used for further footprint analysis. The frequency of footprint colonies was calculated from the number of Amp^R^/Strep^R^ colonies relative to Amp^R^ colonies.

### Detection of excised transposon fragments (ETFs)

HeLa cells cultured at 32°C were transfected with pT_2_-tiHsp70-SB11-SV40-Neo by using the FuGENE^®^ HD Transfection Reagent. Then, cells were incubated at 37°C for 48 hours and selected in medium containing G418 (600 ng/μl) at 32°C. G418-resistant cell colonies were collected and expanded into individual cultures. The G418-resistant cells were heat-shocked for 1, 3, 6, 12, 24 and 48 hours, following by a 2-hour recovery at 32°C. Total DNA was isolated from these cells using a PureLink^®^ Genomic DNA Mini Kit (Invitrogen). Forty μg genomic DNA was used for detection of ETFs. Southern blot hybridization was performed as previously described (38). The DNA probe for *neo* was amplified with two primers (Neo-For and Neo-Rev) and labelled using the DIG high Primer DNA Labeling and Detection Starter Kit II. Hybridization and detection were carried out according to the manufacturer’s instructions.

### 5’-phosphate and 3’-hydroxyl groups of ETF termini

HeLa cells were transfected with pT_2_/tiHsp70-SB11-SV40-Neo (intact transposon), TAN_4_Lo (lack of left flanking TA dinucleotides) or Lo(S3) (mutation of left adjacent CAG to CAA), and selected in medium containing G418 at 32°C. Total DNA was isolated from G418-resistant cells after heat-shock at 37°C for 24 hours following by 2-hour recovery at 32°C. Characterization of 3’-hydroxyl and 5’-phosphate groups of ETFs was performed as previously described (19). Two mg of total DNA was used for sucrose gradient sedimentation. Purified ETFs were digested with *Bsp*TI, which cleaved close to the left ends of the transposons. Then, DNA fragments were separated on a 20% denaturing polyacrylamide gel containing 7M urea, run in TBE, transferred to a Hybond-N^+^ (Millipore) by electro-blotting for 2 hours in TBE at 40 V. DIG-labelled oligonucleotides complementary to the sequences under analysis were used as Southern probes.

The 5’- and 3’-strand-specific markers were synthetic oligonucleotides that have the same sequence and length as potential single-stranded products from transposon termini (Table S1). An equimolar mixture (0.05 pmol/each) of six phosphorylated or unphosphorylated oligonucleotides (24-, 25-, 26-, 27-, 28- and 29-mer), were used as the 5’-phosphorylated or 5’-unphosphorylated markers. An equimolar mixture of seven phosphorylated oligonucleotides (28-, 29-, 30-, 31-, 32-, 33- and 34-mer) was used as the 3’-phosphorylated markers.

### Terminal overhangs of ETFs

Detection of ETF termini overhangs was performed following a protocol as described in Figure 6A. First, pT/SVneo (1 μg) was transfected without (control) or with pCMV-SB11 (200 ng) into 3×10^5^ HeLa cells, or pT/SVneo (100 ng/μl) was microinjected without (control) or with capped SB11 mRNA (20 ng/μl) into one-cell-stage zebrafish embryos. Three days post-transfection or six hours post-microinjection, total DNA was extracted from cells/embryos lysis by using PureLink^®^ Genomic DNA Mini Kit (Invitrogen).

We used splinkerette PCR (39) to detect the terminal overhangs of ETFs. Adaptors (Table S6) were made by heating equimolar amounts of 100 mM primerettes with 100 mM corresponding splinks at 80°C for 5 min and cooling down to room temperature naturally. Adaptors (7.5 μM) were ligated with T4 DNA ligase to total DNA (50 ng/μl) extracted from cells or embryos. Termini of ETFs were amplified with adaptor-mediated ligation PCR. The primary PCR round was performed with primer pairs of primerette-short/long IR/DR(L2) for the left termini, or primerette-short/long IR/DR(R) for the right termini (Table S1). The second PCR round was performed with primer pairs of primerette-nested/newL1 for the left termini or primerette-nested/IR/DR(R)KJC1 for the right termini (Table S1). PCR products, about 162-bp from the left termini and about 229-bp from the right termini, were ligated into the TA-vector for sequencing.

### ETF-capture assay

HeLa cells were transfected with the plasmid pT_2_/tiHsp70-SB11-SV40-Neo at 37°C for 48 hours and selected with G418 at 32°C. G418-resistant cells were then transfected with pT_2_/CMV-Bsd at 37°C for 48 hours, and selected with both G418 and Blasticidin S (Bsd) at 32°C. Double G418 and Bsd resistant cells were either heat-shocked at 37°C (+SB) or maintained at 32°C (-SB) for 24 hours, and then total DNA was extracted. One hundred mg total DNA was used for sucrose gradient sedimentation to enrich ETFs. Enriched ETFs were ligated to the capture vectors that were generated by restriction endonucleases digestion (Tables S2 and S3). Three capture vectors (pT/Mid11, pT/Mid12 and pT/Mid13) were treated with CIP to remove the 5’ phosphates to prevent self-ligation and self-circulation. Each capture vector contained a 10-bp sequence near the restriction endonuclease sites, which could be used as a unique barcode for each individual vector. The ligation mixtures were transformed into competent *E. coli* cells. The double Amp/Bsd resistant colonies were counted and detected with DNA sequencing.

### ETF integration assay

Artificial ETFs were generated by digesting the reconstructive vectors with appropriate restriction endonucleases as described in Tables S4 and S5. The positive control was transfected with pCMV-SB11 and intact plasmid pT_2_/zbHsp70-Bsd, in which the *Bsd* gene was under the control of a zebrafish heat-shock promoter (zbHsp70) (35). One μg purified artificial ETFs were co-transfected with or without pCMV-SB11 (200 ng) into HeLa cells (3×10^5^). Two days after transfection, 1×10^4^ cells were seeded onto 10-cm plates and selected with Bsd (10 ng/μl) for two weeks. Plates with Bsd-resistant cell colonies were fixed with 10% (v/v) formaldehyde in PBS for 15 minutes, stained with 0.1% (w/v) methylene blue in PBS for 30 minutes, washed extensively with deionized water, air dried, counted, and photographed.

Genomic DNA from Bsd-resistant cell colonies was digested with *Sau*3AI and integration sites of ETFs in genomic DNA were analyzed by using the splinkerette PCR method as described previously (39).

## RESULTS

### MMR is not required for resolution of SB transposon excision and formation of canonical and non-canonical footprints

The canonical footprints generated by SB transposase contain three nucleotides from the ETF termini and two TA dinucleotides at each end (Figure. 1B), are proposed to result from the DNA mismatch repair (MMR) process generated by symmetrical cleavages (20). However, this may not be the only pathway that ETFs and donor DNA may take if the symmetry of the excision process is imprecise.

To investigate whether MMR is required for SB footprint formation, cultured mammalian cells were co-transfected with pCMV-SB11 (12) and SB transposon vectors (Figure 1C). We found that the most common footprints were TACAGTA and TACTGTA, as expected (20,36,40), in both HeLa and HCT116 (a colon cancer cell line that is deficient in DNA MMR activity (41)). In both cell lines we found the same ratio of the two canonical footprints with the TACAGTA > TACTGTA (top two lines in Figure 1D). However, we observed that more than half of the footprints detected by this assay could be divided into six non-canonical footprints with flanking TA dinucleotide [TACATA, TACTA, TATGTA, TAGTA, TACGTA, and TATA (which had no base pairs between the flanking TA motifs)] in both HeLa and HCT116 cells (Figure 1D). The canonical model that invokes a 3-bp staggered cut followed by MMR-mediated repair in the formation of SB-mediated footprints does not easily account for these non-canonical sequences that vary.

To explore further the involvement of MMR in footprint formation, we performed plasmid-rescue assays in which we looked for the repair of plasmid backbone by MMR after excision. Figure S2A outlines our experimental approach wherein SB11 transposase is under control of a tiHSP promoter to allow evaluation of excision and repair in MMR-competent cells (HeLa) and MMR-incompetent cells (HCT116). The transposon contains the *rpsL* gene that confers sensitivity to streptomycin (37). Plasmid DNA is recovered from the transfected cells and transformed into *E. coli*. Following transposon excision and repair of the plasmid backbone, the *rpsL* is lost thereby rendering bacteria Strep^R^. As shown in Figure S2B, we found no significant difference between the HCT116 cells [0.11% (378/3.44×10^5^) without heat shock and 0.27% (892/3.30×10^5^) with heat-shock] and the HeLa cells [0.12% (556/4.63×10^5^) without heat shock and 0.33% (1.15×10^3^/3.48×10^5^) with heat shock] in the frequency of Amp^R^/Strep^R^ colonies relative to just Amp^R^ colonies. In support of our findings shown in Figure 1D, we detected similar levels of canonical and non-canonical SB-mediated footprints in the recovered plasmids isolated from the Amp^R^/Strep^R^ colonies in both MMR-competent and MMR-incompetent cells (Figure S2C). These results suggest that MMR is not necessary in footprint formation.

We used the same strategy as that shown in Figure 1C to examine SB footprint formation in NHEJ repair-deficient cell lines XRS5 (42) and M059J (43). We found that the frequencies of SB footprints were diminished while the frequency of deletions was greatly increased (Figure S3). These results suggest that NHEJ activity is very important for excision repair.

### Excision of SB transposon requires at least one flanking TA and adjacent CAG sequences

The footprint patterns shown in Figure 1D can be accounted by asymmetrical cleavage. Asymmetrical cleavage could generate at least twenty-eight ETFs and accompanying footprints, two canonical (TACAGTA and TACTGTA) and six non-canonical (TACATA, TATGTA, TACTA, TAGTA,TACGTA, TATA) footprints (Figure 2). We hypothesize that these ETFs and footprints could arise from non-standard, asymmetrical cleavages at multiple sites within the flanking TA dinucleotide and terminal CAG sequences. To test this hypothesis, we generated model plasmids that either lacked the flanking TA dinucleotide or contained mutated terminal CAG sequences (Figure S1). As shown in Figure 3A, deletion of the left TA (TAN_4_Lo) resulted in the footprint CAGTA that lacks the left flanking TA, and deletion of the right TA (TAN_4_Ro) resulted in footprints TACAG and TACTG that lacked the right flanking TA. In contrast, deletion of both left and right flanking TA dinucleotides (TAN4Lo/TAN4Ro) did not produce any footprints, including either CAG or CTG. These data suggest that SB transposase minimally requires at least one TA site juxtaposed to an outer DR.

**Figure 2.**
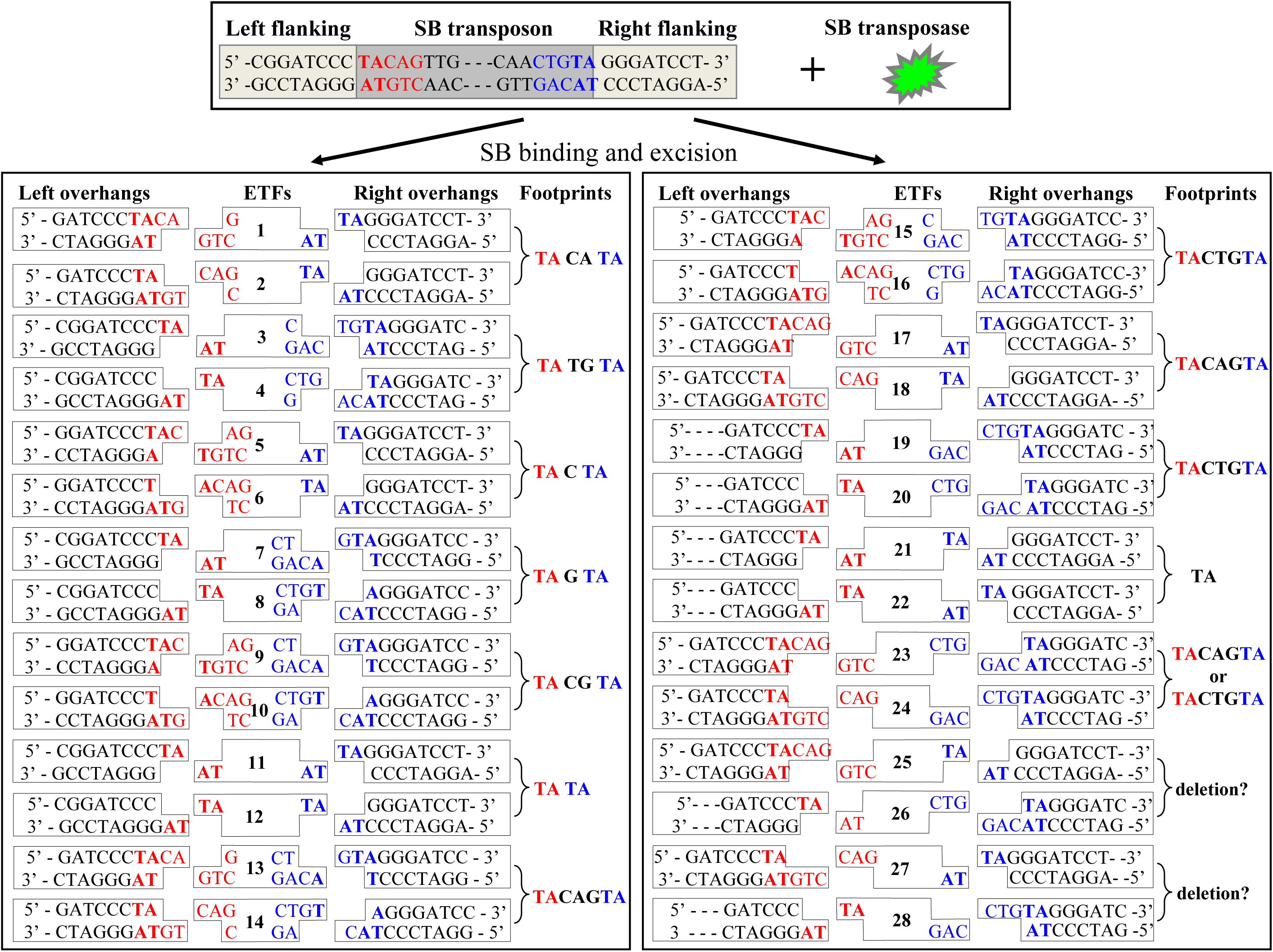
Asymmetrical cleavage products of SB transposase. Multiple possible ETFs and resulting donor DNA structures that contain either 2- or 3-bp overhangs and left two 7-bp footprints (TACAGTA and TACTGTA), six other footprints (TACATA, TATGTA, TACTA, TAGTA, TACGTA, TATA). Portions of the original left and right terminal sequences and two TA dinucleotides are color-coded in the top panel to facilitate the origins of sequence motifs shown in the bottom panel. The potential ETFs are numbered 1-28 in their centers.

**Figure 3.**
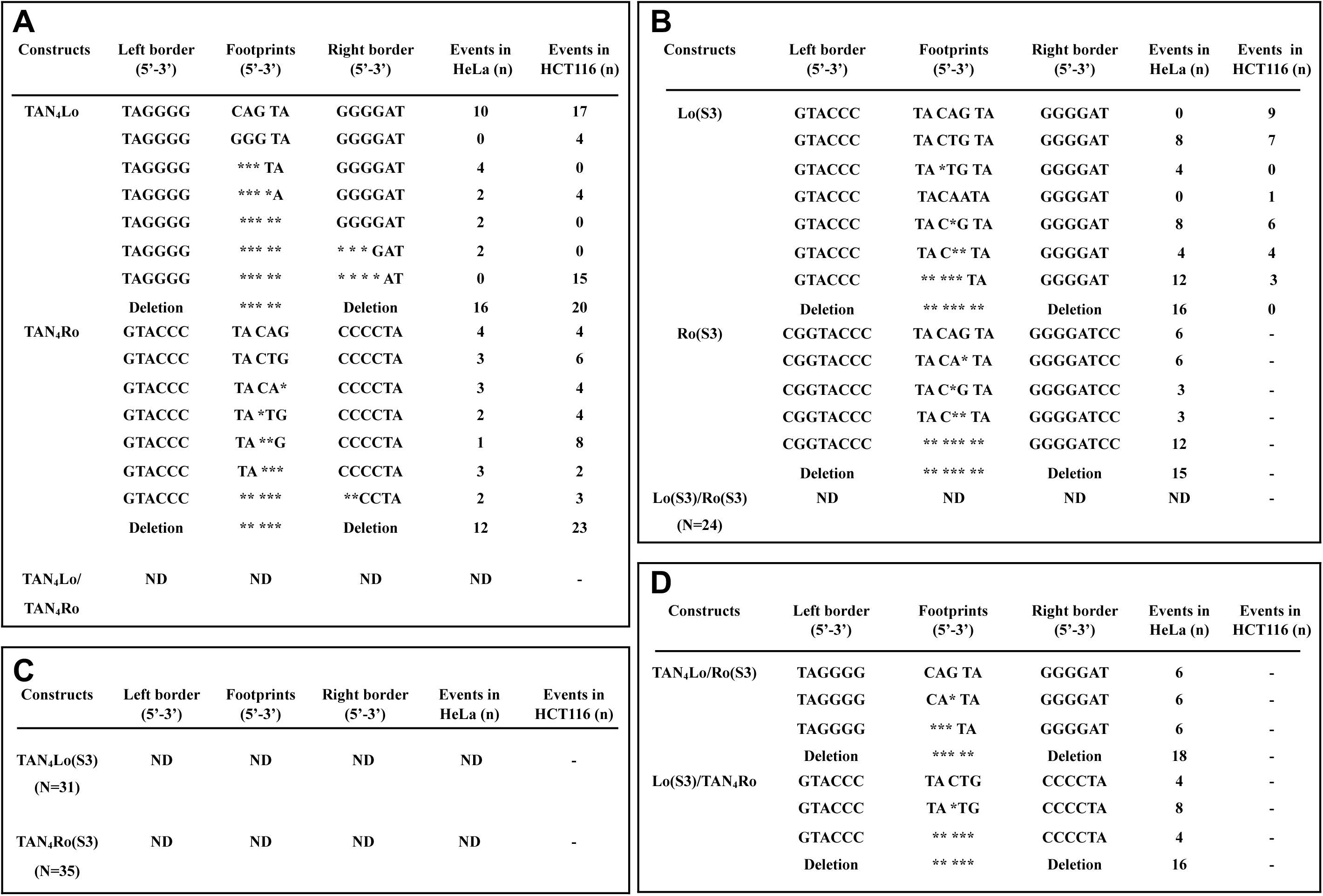
Model SB transposons with altered flanking sequences and the footprints they generate. Variations of canonical SB transposon ITRs and flanking TA dinucleotides (Fig. 1A) were constructed and footprints left in either HeLa (MMR-competent) or HCT116 (MMR-deficient) cells were determined (righthand columns in each panel). The left and right borders of the constructs are shown in the second and fourth from the left columns along with the footprints (third column). * indicates a missing base from canonical footprints; ND, not detectable; -, not tested. Data were pooled from six independent experiments in each of the two cell lines. (A) Effects of deleting the TA dinucleotide adjacent to either Lo (TAN_4_Lo), Ro (TAN_4_Ro) or both TAN_4_Lo/TAN_4_/Ro. (B) Effects of converting the CAG sequence to CAA at the terminus of either Lo (S3) or Ro (S3) or both Lo(S3)/Ro(S3). (C) Effects of both deleting the TA dinucleotide and mutating the CAG sequence to CAA at Lo, Ro, or both. (D) Effects of either deleting the TA dinucleotide or mutating the CAG sequence to CAA at Lo and the complementary deletion or mutation of CAG to CAA at Ro.

We next tested the influence of the adjacent CAG sequences on footprint formation. Alteration of CAG to CAA in either the left [Lo(S3)] or right [Ro(S3)] terminus generated the two canonical footprints, as well as other footprints (Figure 3B). However, we could not detect any footprints after mutation at both terminal ends [Lo(S3)/Ro(S3); Figure 3B, bottom line)]. Likewise, deletion of either flanking TA dinucleotide along with alteration of its adjacent CAG to CAA [TAN_4_Lo(S3) and TAN_4_Ro(S3)] resulted in no detectable footprints (Figure 3C). However, deletion of the flanking TA dinucleotide at one end and mutation of the terminal CAG to CAA at the other end [TAN_4_Lo/Ro(S3) and Lo(S3)/TAN_4_Ro] could generate footprints CAGTA and TACTG, respectively (Figure 3D). Taken together, these data indicate that at least one flanking TA dinucleotide and one CAG sequence are required for footprint formation.

### SB transposase generates ETFs of variable lengths

Similar to other Tc1/*mariner* transposons, SB transposition is characterized as a “cut-and-paste” mechanism. However, there is only circumstantial evidence showing the existence of ETFs during transposition by SB (21) and *Hsmar1* (25). To test the asymmetrical excision model in Figure 2, we first looked at the spectrum of ETFs. HeLa cells were transfected with plasmids pT_2_/tiHsp70-SB11-SV40-Neo and selected with G418 (Figure 4A). Southern blots of total DNA isolated from heat-shocked G418-resistant cells, showed the existence of extrachromosomal ETFs that were consistent in size with excised transposons from plasmids (Figure 4B).

**Figure 4.**
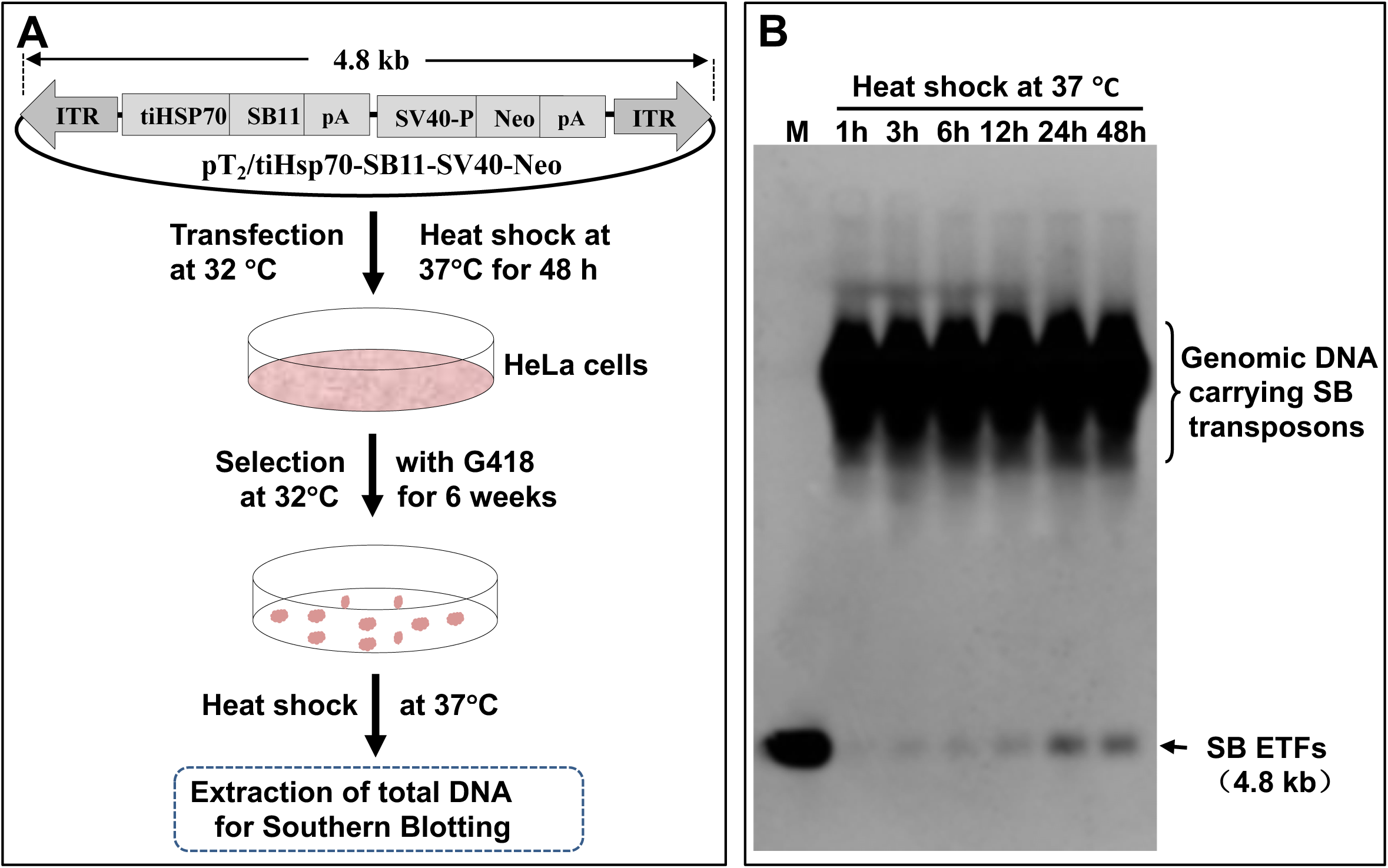
Detection of ETFs. (**A**) Experimental design for ETF detection. HeLa cells cultured at 32°C were transfected with plasmids pT_2_/tiHsp70-SB11-SV40-Neo, in which expression of SB11 transposase is induced by heat-shock at 37°C of a tilapia heat-shock promoter 70 (tiHsp70). The G418-resistance gene (Neo) is controlled by the SV40 promoter (SV40-P), to obtain stably G418-resistant cells at 32°C. (**B**) G418-resistant HeLa cells were heat-shocked at 37°C for 1, 3, 6, 12, 24 and 48 hours and then recovered at 32°C for 2 hours. Total DNA was used for Southern blotting analysis to detect ETFs. M, linear transposons were used as a positive control.

The asymmetrical cleavage model predicts that there will be multiple cleavage sites in an ensemble of excised transposons. Hence, we explored the terminal structures of the ETFs from above. Purified ETFs were digested with *Bsp*TI, which cleaved close to the transposon termini (Figure 5A). The 5’-strands of *Bsp*TI-fragments (5’T_2_) from intact transposons, migrated with the same patterns as those of six phosphorylated synthetic markers (labeled 5’P; Figure 5B). The 3’-strands of *Bsp*TI-fragments (3’T_2_) distributed into six bands similar to six of seven phosphorylated synthetic markers (labeled 3’; Figure 5C). These data support an asymmetrical cleavage model and suggest that SB transposase cleavage positions might be within the flanking TA dinucleotide and adjacent CAG sequences. Likewise, we found that the deletion of left flanking TA dinucleotide led to only the 24-, 25- and 27-mer bands [5’T_2_(,0, TA), Figures 5D and S4] and the 28-, 29- and 31-mer bands [3’T_2_(,0, TA), Figures 5E and S4]. Mutation of the left adjacent CAG to CAA [5’T_2_(G/A) and 3’T_2_(G/A)], resulted in only five bands (Figures 5F, 5G and S4). These data reinforce asymmetric excision model that allows cleavage at multiple sites within the flanking TA dinucleotide and adjacent CAG sequences in the outer DRs.

**Figure 5.**
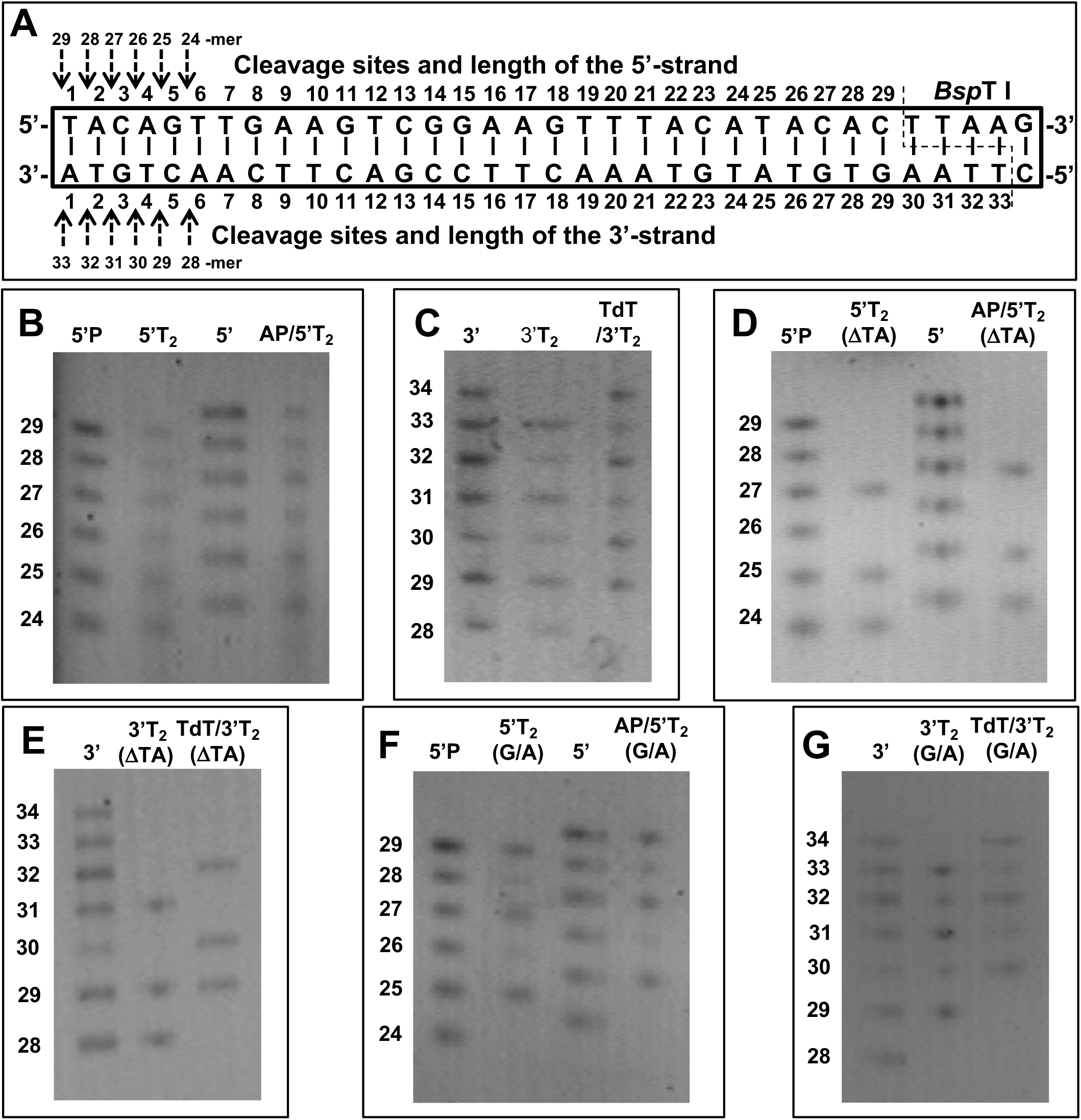
Variation in cleavage-sites by SB transposase. (**A**) The left terminal sequence (Lo) of SB transposon with nucleotides labelled 1-29 (top strand) and 1-33 (bottom strand). The *Bsp*TI site is on the extreme right. Potential cleavage sites on the 5’- and 3’-strands are indicated by dotted arrows with the lengths of the resulting single-stranded sequences shown elevated or lowered on the left side of the panel. The left most lanes in Panels (B) – (G) are markers. 5’P and 5’: phosphorylated and unphosphorylated 24-, 25-, 26-, 27-, 28- and 29-mer synthetic oligonucleotides, which had the same sequence as the 5’-end of ETFs, respectively. 3’: Phosphorylated 28-, 29-, 30-, 31-, 32-, 33- and 34-mer oligonucleotides. (**B**) Excision sites in the 5’-strand of ETFs. 5’T_2_: ETFs digested with *Bsp*TI. AP/5’T_2_: ETFs treated with alkaline phosphatase (AP) to determine the presence of 5’-phosphate groups. (**C**) Excision sites within the 3’-strand of ETFs. 3’T_2_: ETFs digested with *Bsp*TI. TdT/3’T_2_: ETFs treated with ddATP and TdT before digestion with *Bsp*TI. (**D**) Excision sites within the 5’-strand of ETFs lacking the left-flanking TA. 5’T_2_(ΔTA): ETFs lacking the left-flanking TA digested with *Bsp*T I. AP/5’T_2_ (ΔTA): ETFs lacking the left-flanking TA were treated with AP. (**E**) Excision sites within the 3’-strand of ETFs lacking the left-flanking TA. 3’T_2_(ΔTA): ETFs lacking the left-flanking TA digested with *Bsp*TI. TdT/3’T_2_(ΔTA: ETFs lacking the left-flanking TA treated with ddATP and TdT before digestion with *Bsp*TI. (**F**) Excision sites within the 5’-strand of ETFs containing a CAG to CAA mutation. 5’T_2_(G/A): ETFs containing a CAG to CAA mutation digested with *Bsp*TI. AP/5’T_2_(G/A): ETFs containing a CAG to CAA mutation treated with AP. (**G**) Excision sites within the 3’-strand of ETFs containing a CAG to CAA mutation. 3’T_2_(G/A): ETFs containing a CAG to CAA mutation digested with *Bsp*T I. TdT/3’T_2_(G/A): ETFs containing a CAG to CAA mutation treated with ddATP and TdT before digestion with *Bsp*TI.

**Figure 6.**
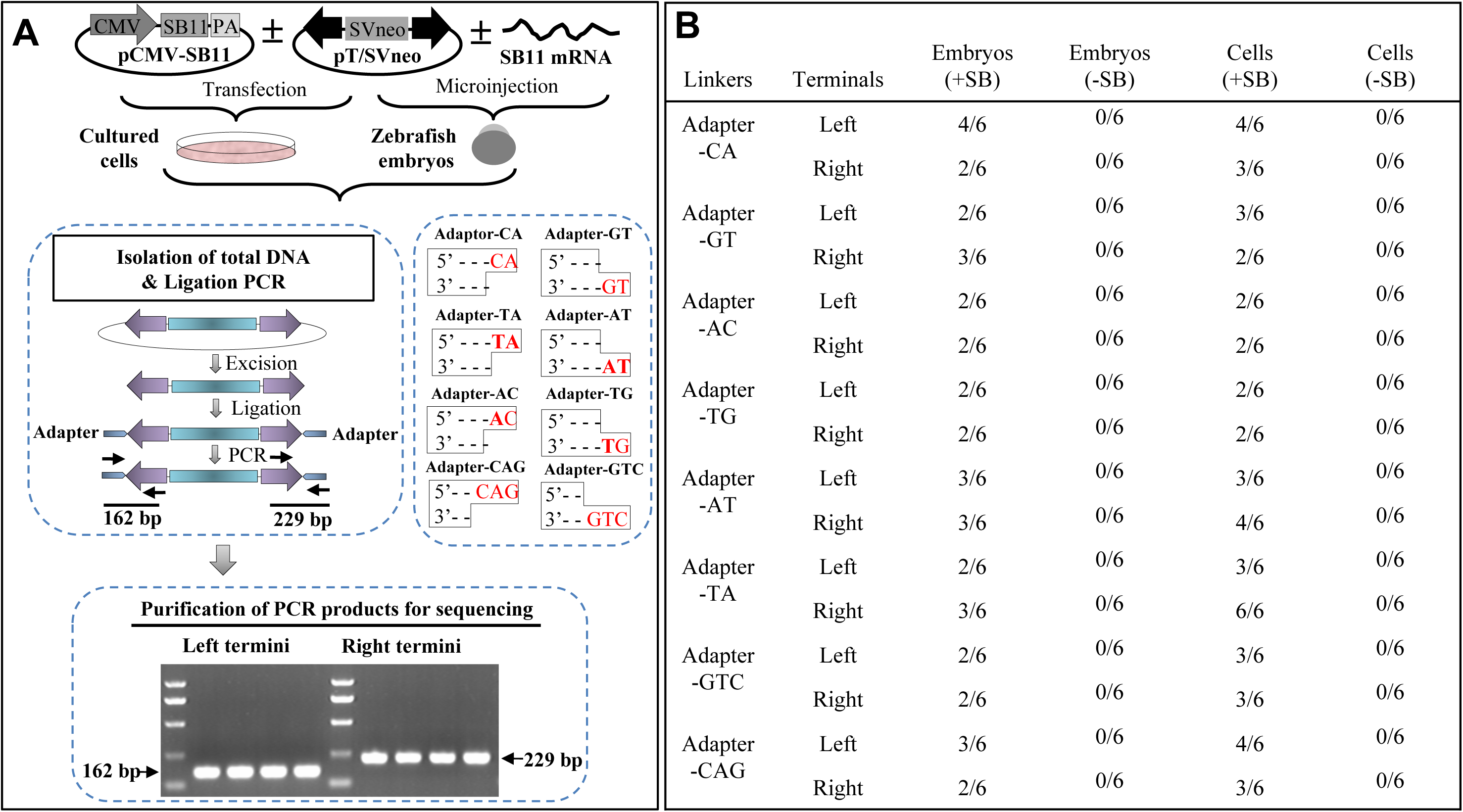
Terminal overhangs of ETFs. (**A**) Experimental design for detection of ETF termini. HeLa cells were transfected with pT/SVneo ± pCMV-SB11 (+SB and –SB). Zebrafish embryos were microinjected with pT/SVneo ± SB11 mRNA (+SB and –SB) and collected at 6 hours post-fertilization. Total DNA from either HeLa cells or embryos was extracted and ligated to appropriate adaptors. Adaptor-mediated PCR was performed to generate products about 162 bp or 229 bp from the left or right termini for sequencing, respectively. (**B**) The frequencies of PCR products containing the precise terminal sequence from six independent experiments are tabulated; no PCR products were detected without heat-shock induction of SB transposase expression.

We also tested whether the 5’-temini of ETFs contained 5’-phosphate groups. We found that alkaline phosphatase (AP) treatment shifted the 5’-strands of *Bsp*TI-fragments to positions corresponding to unphosphorylated markers (right hand columns in Figures 5B, 5D and 5F). These results suggest that the 5’-termini of ETFs were phosphorylated. Likewise, we examined whether the 3’-termini of ETFs possessed hydroxyl groups. Terminal deoxynucleotidyl transferase (TdT) treatment adds extra nucleotides to 3’-ends that have hydroxyl groups. As shown in columns labelled TdT, TdT-treatment with ddATP increased the lengths by a single nucleotide of the *Bsp*TI fragments from either intact transposons (Figure 5C) or reconstructed transposons (Figures 5E and 5G). These data indicate that the termini of ETFs contained 3’-hydroxyl groups.

### Effective integration of ETFs requires a TA and a CAG sequences

To further test the asymmetric excision model, we interrogated the terminal structures of the ETFs. Adaptors carrying the same overhangs as the proposed ETFs were generated (Table S6). As shown in Figure 6A, adaptors were ligated to total DNAs extracted from either HeLa cells or zebrafish embryos. The termini of the ETFs were amplified and the resulting PCR products were sequenced. We detected all the eight overhangs in HeLa cells and zebrafish embryos (Figure 6B) as well as in NHEJ-deficient cells XRS5 (except adaptor-AC) and M059J (Figure S5) in a SB-dependent manner; no PCR products were ever detected in cells or zebrafish embryos without a source of SB. These data support the existence of multiple ETFs following SB-mediated excision.

We next designed a strategy to capture ETFs during transposition (Figure 7A). As shown in Figure 7B, twenty ETF were detected in cells following heat-shock to induce SB expression (ETF1 - ETF20; Figure 2). We can not rule out the presence of other ETFs, including ETF-23 that is proposed as a product of the standard excision model (20). However, if produced, proposed ETFs (ETF21 - ETF28; Figure 2) were below minimal detectable levels.

**Figure 7.**
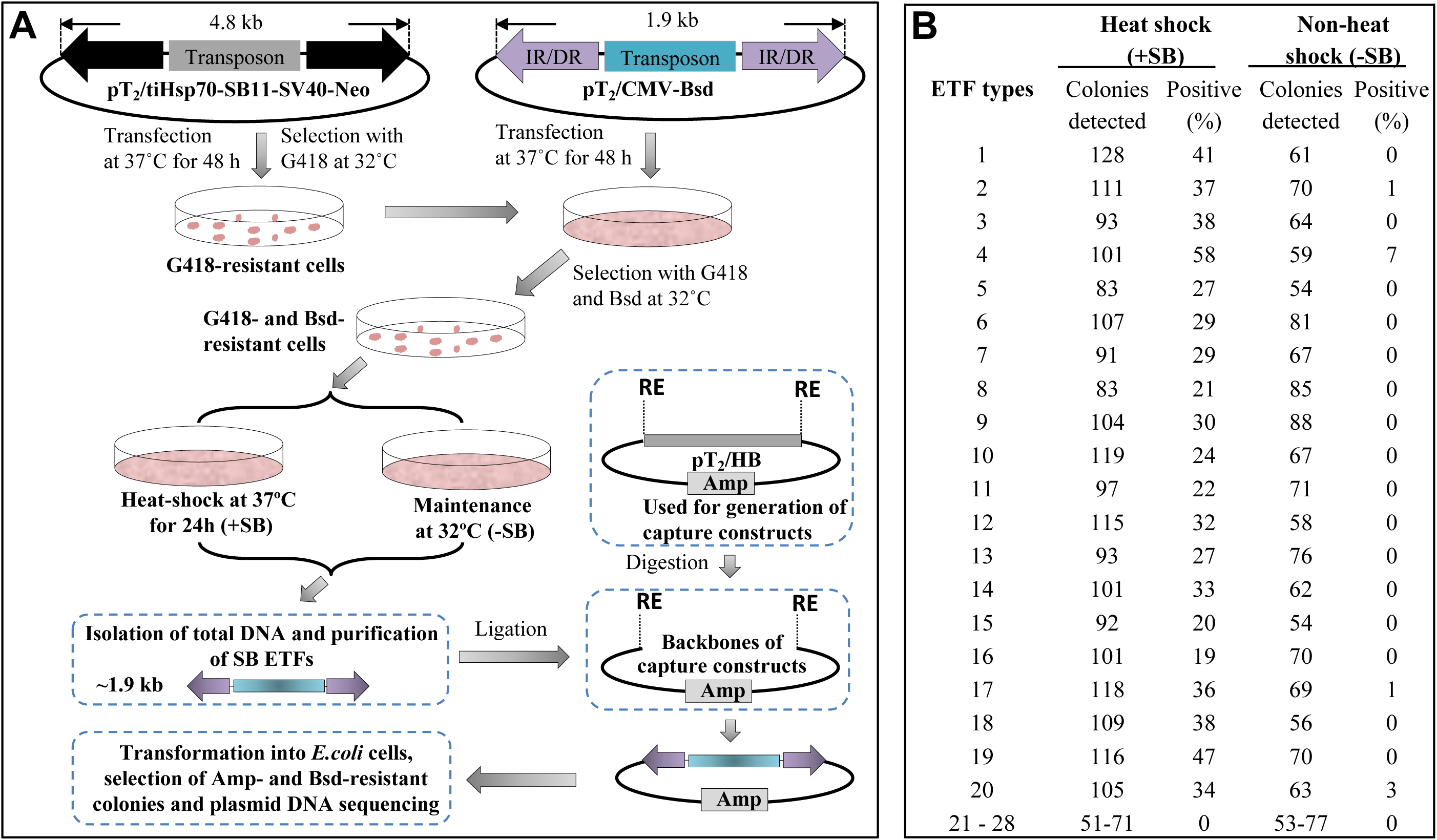
Capture of ETFs. (**A**) Experimental design for ETF detection. G418-resistant cells were transfected with pT_2_/CMV-Bsd-SV40 and selected in Bsd-containing medium. G418- and Bsd-resistant cells were heat-shocked at 37°C to induce the excision of SB transposons under the control of a tilapia heat-shock promoter (tiHsp) promoter and the control cells were maintained at 32°C. Total DNA was extracted and used for purification of ETFs. Purified-ETFs were ligated with sixteen ETF-capturing constructs shown in Table S3 and transfected into Top10-competent *E. coli*. (**B**) ETF sequences from Amp- and Bsd- resistant colonies from (**A**) were amplified by PCR and sequenced to quantify relative levels of ETFs generated following induction of SB expression. Data were pooled from three independent experiments.

To test whether the detected ETFs could effectively integrate into a genome, we designed constructs that could generate the twenty-eight ETFs shown in Figure 2. Transposition assays were performed by co-transfection of artificial ETFs with or without pCMV-SB11 into HeLa cells (Figure 8A). As shown in Figures 8B and S6, plasmids with a standard sequence showed the highest transposition rates (100%) followed by eight ETFs (ETF1 - ETF4 and ETF17 - ETF20). The other twelve ETFs (ETF5 - ETF16) and the eight undetectable ETFs (ETF21 - ETF28) did not produce colony numbers above background. Cell colonies from the eight most prominent ETFs (ETF1 - ETF4 and ETF17 - ETF20) were expanded for analysis of integration sites in the genome. As shown in Figure 8C, these eight ETFs could integrate into TA loci as expected. In sum, eight of the twenty-eight theoretical ETFs were able to effectively complete the integration process during SB transposition and these ETFs had at least one TA dinucleotide and a single-stranded portion of the CAG sequence.

**Figure 8.**
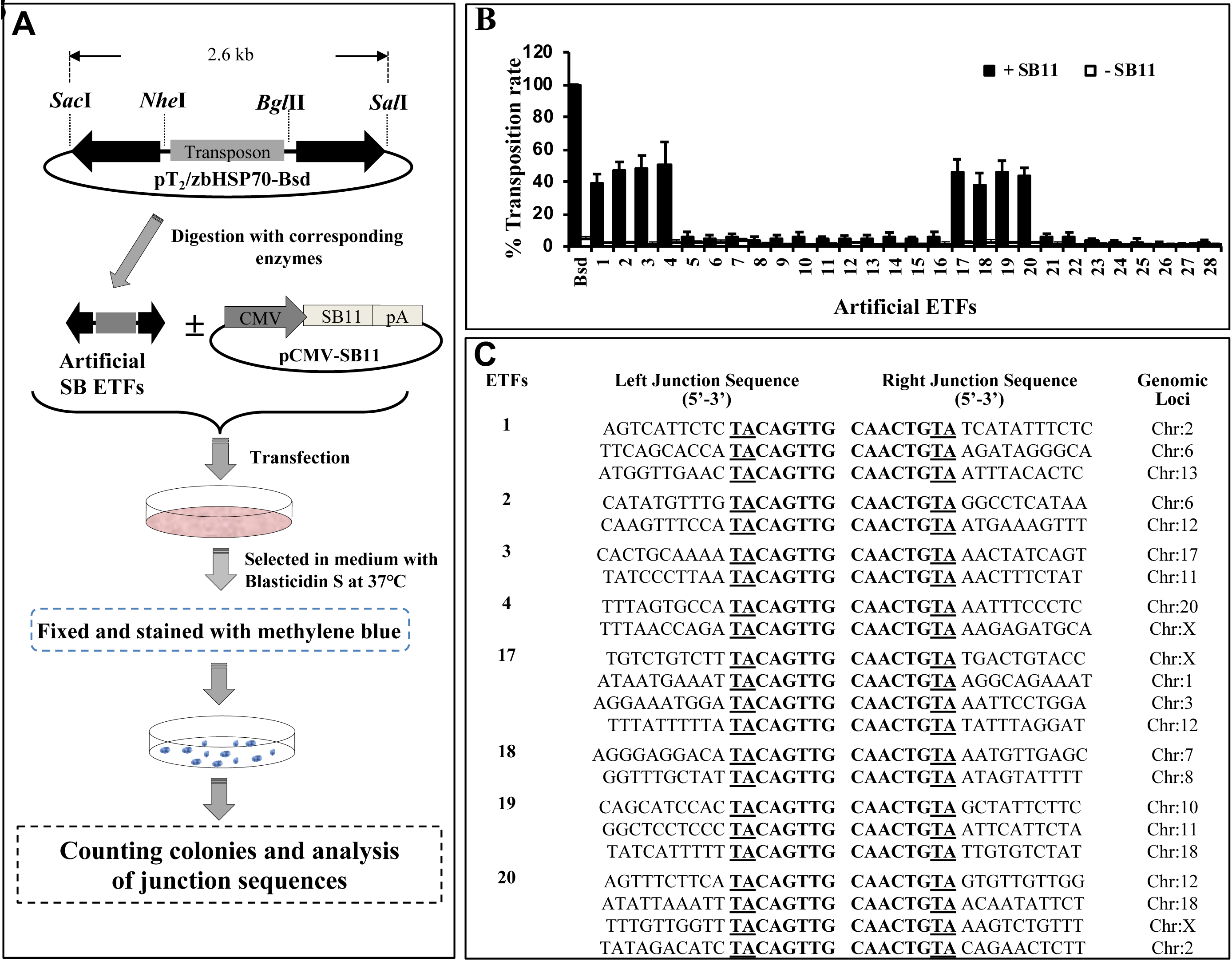
Transposition efficiencies of model ETFs. (**A**) pT_2_/zbHsp70-Bsd plasmids were modified to mimic twenty-eight ETF sequences representing the those shown in Figure 2. Artificial ETFs were co-transfected with or without pCMV-SB11 into HeLa cells and selected with Bsd at 37°C for about two weeks. The Bsd-resistant cell colonies were fixed with formaldehyde, stained with methylene blue and counted, and photographed. (**B**) Transposition rates of the model ETF transposons were calculated by the number of cell colonies from three independent experiments. (**C**) Genomic DNAs of individual colonies from model ETF transposons (ETF1 - ETF4 and ETF17 - ETF20) were isolated for integration site analysis. The flanking TA dinucleotides of SB transposons are underlined.

## DISCUSSION

### Roles of TA dinucleotides and adjacent CAG sequences in SB transposition

Our previous studies showed that having a TA-dinucleotide flanking one side of a SB transposon is essential for excision and reintegration and that substitutions of the first or third base in the adjacent CAG markedly reduced the activities of excision and transposition. In contrast, mutation of both the first and third base in the CAG led to undetectable activities of excision and transposition (7, 36). Therefore, the TACAG sequence at the end of SB transposon is essential for excision and transposition.

A previous study has shown that two ITRs of SB transposon are functionally not equivalent since the left ITR contains sequences required for high efficiency transposition (44). Recently, aberrant transposition events, including single-ended transposition of *Mos1* and *mariner* elements, have been observed under suboptimal conditions (21). However, it is unclear how the TA dinucleotide and its adjacent CAG sequence were involved in transposition. In this study, we found that effective excision and footprint formation required that one terminus of an SB transposon be flanked with a TA dinucleotide and the other have an intact CAG or CTG sequence (Figures 1D and 3). These findings did not support the excision models proposed for Tc3 (45) and SB (1, 20) in which the TA sites are not involved through staggered cuts. Indeed, each nucleotide within the TA and its adjacent CAG appears to be cleavable by SB transposase (Figures 5B and 5C) to generate at least twenty extrachromosomal ETFs (Figures 2, 4 and 7).

Excised but not reintegrated molecules were observed in Tc1 and Tc3 transposition (45, 46). There has been debate whether these extrachromosomal transposon-derived molecules are *bona fide* ETFs or side products of a repair process. For instance, SB-K248A generated extrachromosomal circles that integrate poorly and are neither considered as ETFs nor natural transposition products (8). In this study, ETFs that contained the TA dinucleotide on one terminus and the CAG sequence on the other end were active for integration. In contrast, other ETFs that did not contain either a TA and CAG on each end were ineffective for integration (Figure 8). The failure to capture ETF21 – ETF28 (Figure 7B) and the appearance of few positive cell colonies for integration (ETF21 - ETF28; Figure 8B) suggest that these ETFs either were not formed or not detectable products of SB excision. Thus, re-integration by SB transposase appears to accommodate asymmetrical cleavage of both the flanking TA dinucleotide and adjacent CAG sequences.

A previous study reports that excision efficiency and re-integration rates of SB transposition are not coupled (47). Our capture assays showed the presence of twenty ETFs after SB excision (Figure 7B). These ETFs do not appear to be dependent on an opening of a hairpin structure potentially present on transposon termini. A hairpin-opening activity is not essential in all transpositions because *Mos1* does not proceed through hairpin ETFs (18) and Artemis, which is required for opening the hairpins, is dispensable for SB transposition (48). Accordingly, in the absence of hairpin ETFs, *mariner* and SB transposases may cleave two strands of the DNA at each end of a transposon by two hydrolysis reactions (5) within the DNA/transposase complex that contains the target DNA.

### Repair of SB transposase-mediated excision does not require MMR activity

MMR is a process that recognizes and repairs erroneous insertions, deletions or mis-incorporations of bases during DNA replication and recombination as well as mismatches that result from some forms of DNA damage (49). MMR is proposed to participate the formation of footprints at the donor sites after transposon remobilization (20, 45). The G-G or T-G mismatch can be effectively repaired in HeLa cells, but not in HCT116 cells that have a defective hMLH gene (41). Nevertheless, SB footprints in HCT116 cells were similar to those in HeLa cells (Figure 1D). Similarly, footprints in rescued plasmids from HCT116 (MMR*^-^*) cells were similar in proportion to those in HeLa (MMR*^+^*) cells (Figure S2C). These data suggest that MMR does not play a crucial role in the SB footprint formation. Rather, NHEJ appears to be mainly responsible for the formation of multiple SB footprints through direct ligation of two single-stranded donor ends followed by gap DNA repair. NHEJ factors including Ku, DNA-PKcs and Xrcc4, are required for efficient SB-mediated transposition and repair of excision sites in somatic cells (40, 48). In the absence of either Ku or ATM, a homology-dependent repair pathway and synthesis-dependent strand annealing (SDSA) pathway appear to be initiated (48). In this study, an increased number of deletions were detected at the donor sites of rescued plasmids isolated from two NHEJ-deficient cells (Figure S3), suggesting that NHEJ plays an important role in the formation of SB footprints. However, we can not exclude the involvement of other DNA-repair associated pathways and processes in the high-fidelity DSB repair process following SB excision, because four common footprints (TACAGTA, TACTGTA, TATGTA, TACTA) were detectable in NHEJ-deficient cells (Figure S3). These findings are consistent with the observation that the repair of transposase-induced DSBs in mouse liver is rapid and efficient and does not require the activity of the DNA-PK complex (40).

Multiple lines of evidence from this study indicate that the common footprints result from the repair of SB-mediated excision at the donor sites, rather than from endonuclease activity. First, we detected six 5’-strand bands (5’T_2_) and six 3’-strand bands (3’T_2_) following digestion of purified ETFs with *Bsp*TI (Figure 5). The deletion of the left flanking TA dinucleotides led to three 5’-strand [5’T_2_(LTA)] (Figure 5D) and three 3’-strand bands [3’T_2_(LTA)] (Figure 5E). In addition, mutation of left adjacent CAG to CAA [5’T_2_(G/A) and 3’T_2_(G/A)], resulted in five 5’-strand and 3’-strand bands (Figures 5F and 5G), respectively. The modification of excised overhangs by endonuclease activity does not account for these specific changes. Second, eight ETF overhangs were detectable in HeLa cells and zebrafish upon the induction of SB transposase expression (Figure 6B), indicating that what we observe is neither cell-type dependent nor species dependent. Third, twenty ETFs were detectable in a SB expression-dependent manner (Figure 7B). Importantly, only eight of twenty validated ETFs could efficiently reintegrate to complete transposition (Figures 8B and S6). In totality, these findings support the conclusion that the variable ETFs observed in the several experimental approaches we employed were generated by SB catalysis rather than endonuclease degradation.

Previous studies (20,36,40,48,50) and this study (Figure 1D) have identified a large number of alternative footprints. Similarly, numerous other footprints have also been found in the genomes of transgenic lines after remobilization of *Tc3* elements in addition to the two most common footprints TACATA and TATGTA (45). Based on our SB excision data, eight validated ETFs (ETF13 - ETF20; Figure 2) were responsible for the generation of the two most common 7-bp footprints. Interestingly, SB ETFs 1-4 gave rise to the two common footprints TACATA and TATGTA, which mimic the two most common Tc3 footprints. One of the two most common SB footprints, TACTGTA, has been detected in Tc3 transposition (45). Further investigation of the structural and biochemical activity differences of SB and Tc3 and other transposases would offer opportunities to understand better the detailed events of selection of binding sites that lead to effective excision.

### Implications of our refinements in the standard model of SB transposition

Based on data here and elsewhere, we propose the following refinements of the canonical model for SB-mediated transposition (Figure 9). The process of SB transposition can be divided into six sequential steps: i) Binding of SB transposases to each ITR (51). ii) Formation of a DNA/transposase synaptic complex (5). iii) Asymmetrical cleavages of transposons to generate at least twenty ETFs, and parallel repair of the donor sites by NHEJ thereby leaving eight common footprints (TACTGTA, TACAGTA, TACATA, TACGTA, TATGTA, TACTA, TAGT, TATA). iv) Recognition and cleavage at a TA dinucleotide by the paired-end DNA/transposase complex. v) Integration of the ETF into target TA. vi) Repair of DNA gaps within the integration sequence that leads to a duplication of the original TA sequence.

**Figure 9.**
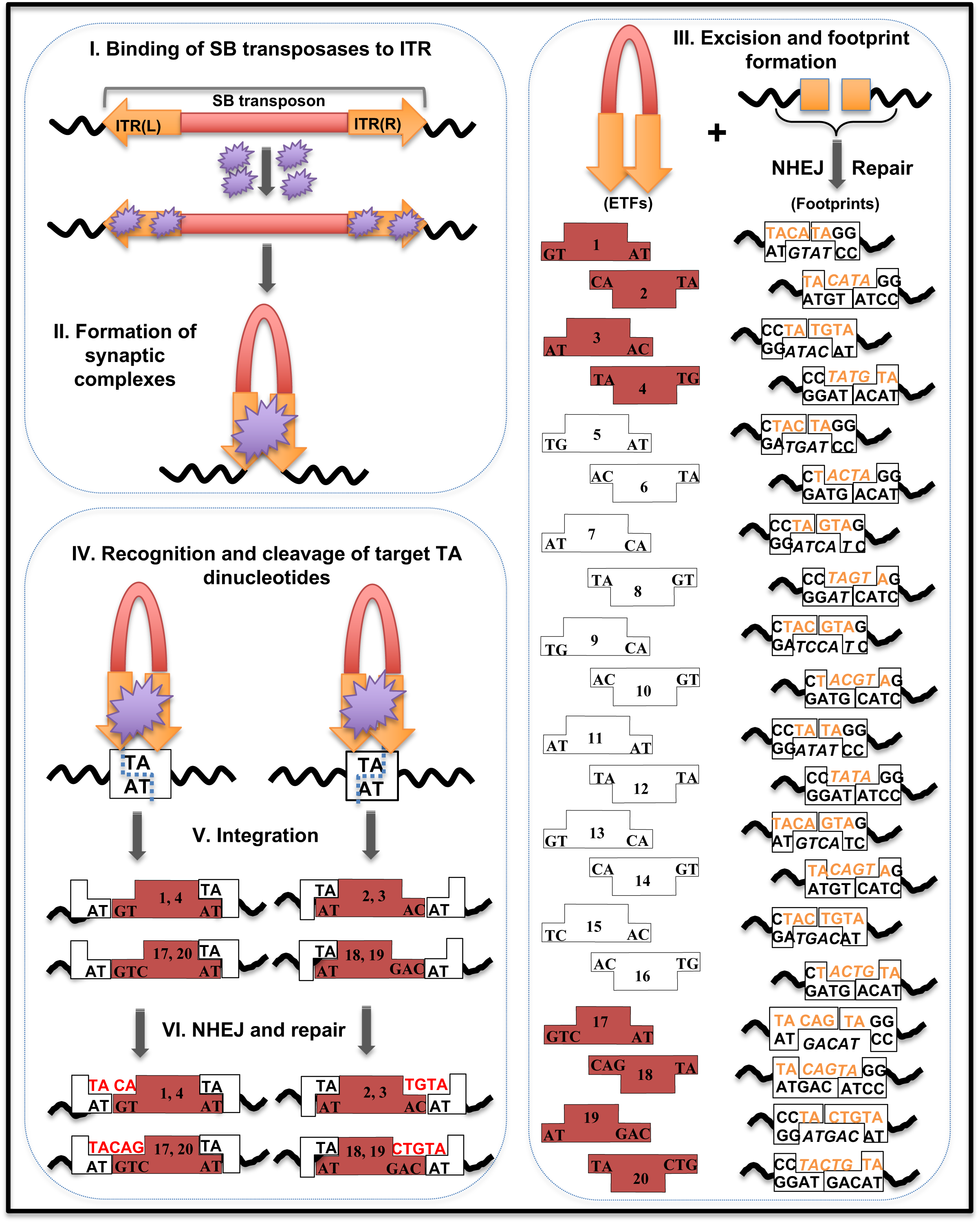
Refined model for SB-mediated transposition. SB transposition can be divided into six sequential steps. In this model, only eight ETFs, ETF1 - ETF4 and ETF17 - ETF20 shown in red, each of which contains one 5’-TA or 3’-AT overhang at one terminus and one 2-bp or 3-bp overhang at the other terminus, can potentially reintegrate into a target TA dinucleotide within the genomic DNA. Since the biochemical structure of an active SB transpososome remains uncharacterized, one large SB transposase complex representing the dimerization (or tetramerization) of transposases binding to inner and outer DRs is shown.

This refined model accounts for the following observations and predictions: 1) The presence of multiple SB-mediated ETFs (48, 52) that may be generated through asymmetric cleavage of transposon termini. 2) Rates of excision and reintegration are not equal (21,47,53,54). Many ETFs are not integrated at detectable levels although there does appear to be a pretty constant ratio of integration/excision that might allow the prediction of a rough level of integration events based on determination of excision products. 3) NHEJ participation in both footprint formation and reintegration may vary to a greater extent than previously thought. 4) MMR is not necessary for SB transposition. In addition, our data may inform on the dynamics of multimeric assembly of SB transposases in the synaptic complex. Selective mutations in the outer DRs suggest that anchoring of SB transposase at the outer DRs is essential for cleavage and that it cannot rely on dimerization (or tetramerization) of transposases binding to inner DRs. This teasing observation exacerbates the question of the mystery roles of the inner DRs that are essential (7) for SB- mediated transposition.

## ACKNOWLEDGEMENT

We thank all members in Cui’s laboratory and Dr. Yuan Xiao and Yan Wang of the Analysis and Testing Center at IHB for technical supports.

## SUPPLEMENTAL INFORMATION

### Supplemental Figure Legends

**Figure S1.**
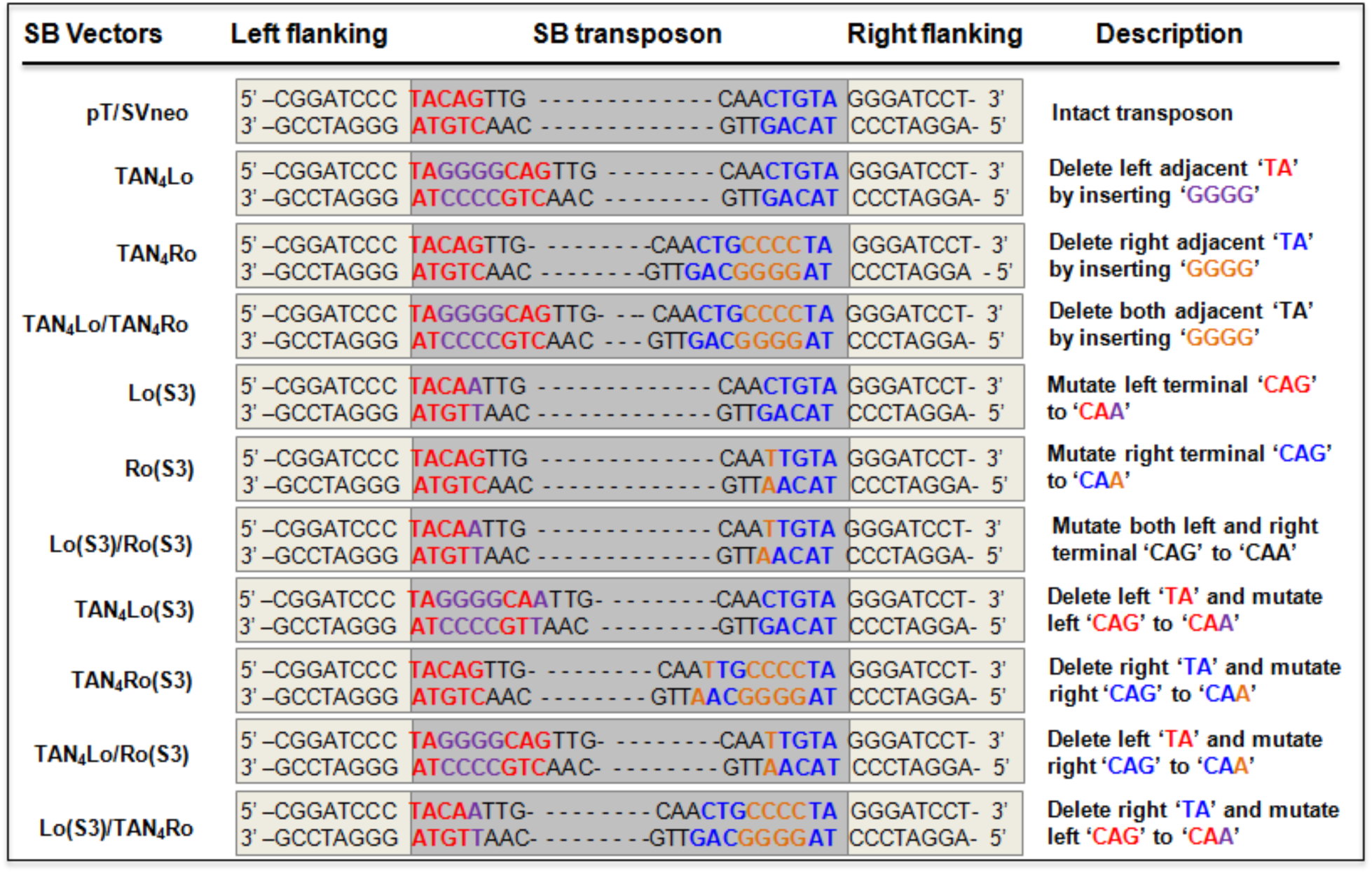
Reconstructive SB transposons. Sequences and descriptions of left and right terminals of different SB transposons.

**Figure S2.**
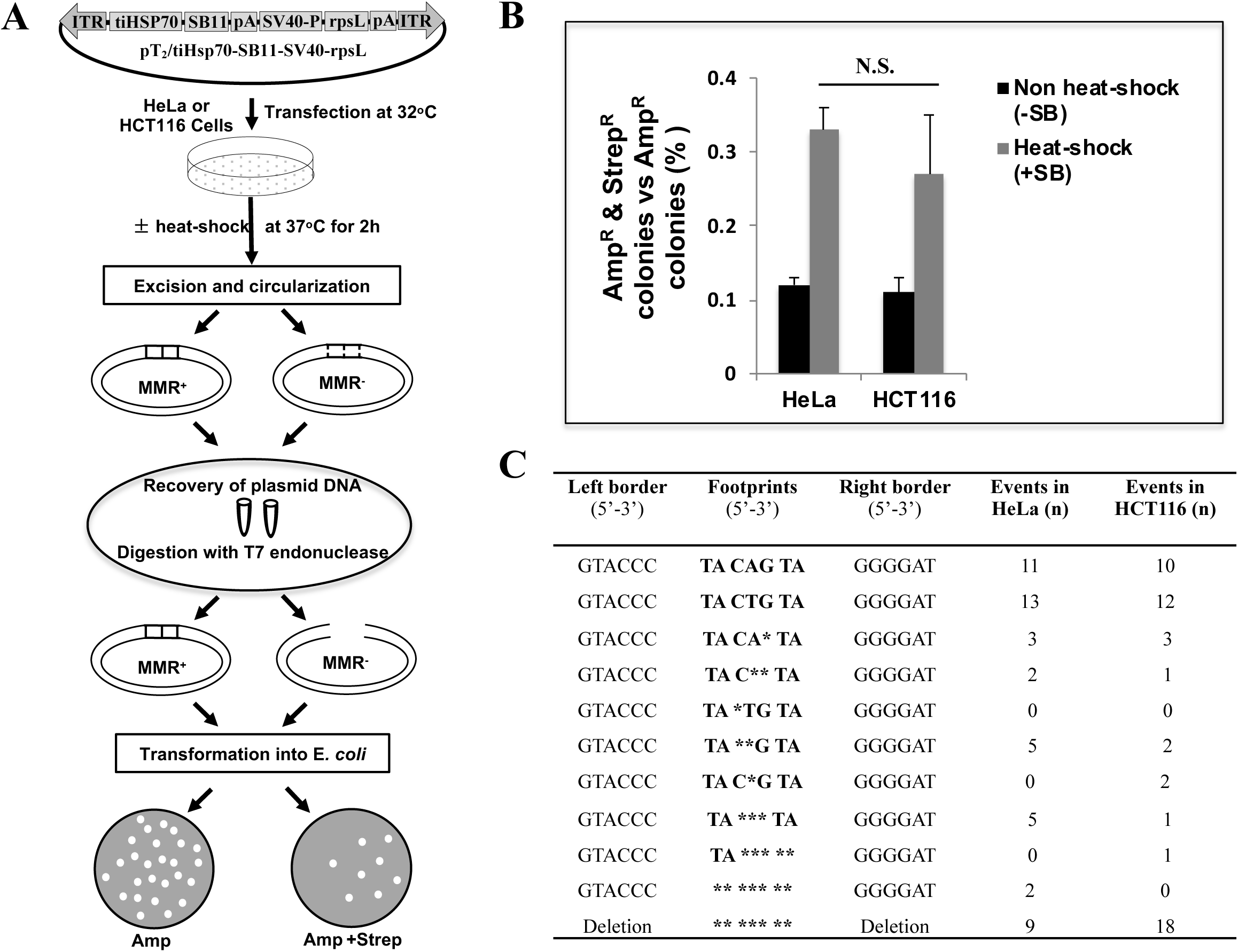
Roles of MMR in the formation of SB footprints. (A) The details of plasmid rescue assay were described in the materials and methods. Briefly, Plasmid DNA was transfected into HeLa (MMR^+^) and HCT116 (MMR^-^) cells and cultured at 32°C. Twenty-four hours after transfection, cells were either induced at 37 °C (heat-shock) for 2 hours or maintained at 32 °C (non heat-shock). Plasmid DNA recovered from transfected cells was subjected to T7 endonuclease I (T7) digestion. Digested DNA products were then transformed into competent Top10 *E.coli.* cells. Transformed *E. coli*. cells were then subjected to a selection of either Amp (100 μg/ml) or a double selection of Amp/Strep (100 μg/ml and 30 μg/ml). tiHSP70, tilapia HSP promoter; SV40 P, simian virus 40 promoter; pA, poly(A). **(B)** Amp^R^/Strep^R^-bacteria colonies were named as footprint colonies, and frequency of footprint colonies was calculated as Amp^R^/Strep^R^ normalized by Amp^R^. Data were pooled from six independent experiments in each of the two cell lines.**(C)** Amp^R^/Strep^R^-bacteria colonies were used for footprint analysis.

**Figure S3.**
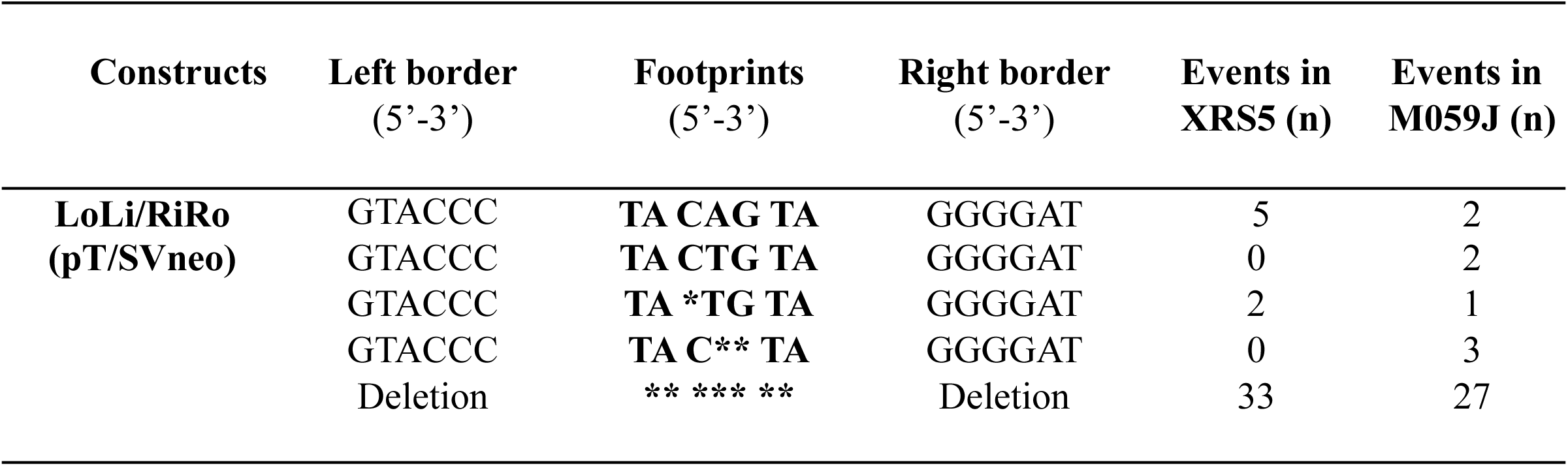
SB footprints in NHEJ deficiency cells. The XRS5 and M059J cells were transfected with pCMV-SB11 and pT/SVneo. Footprints were detected. **\*** indicates a missing base from canonical footprints.

**Figure S4.**
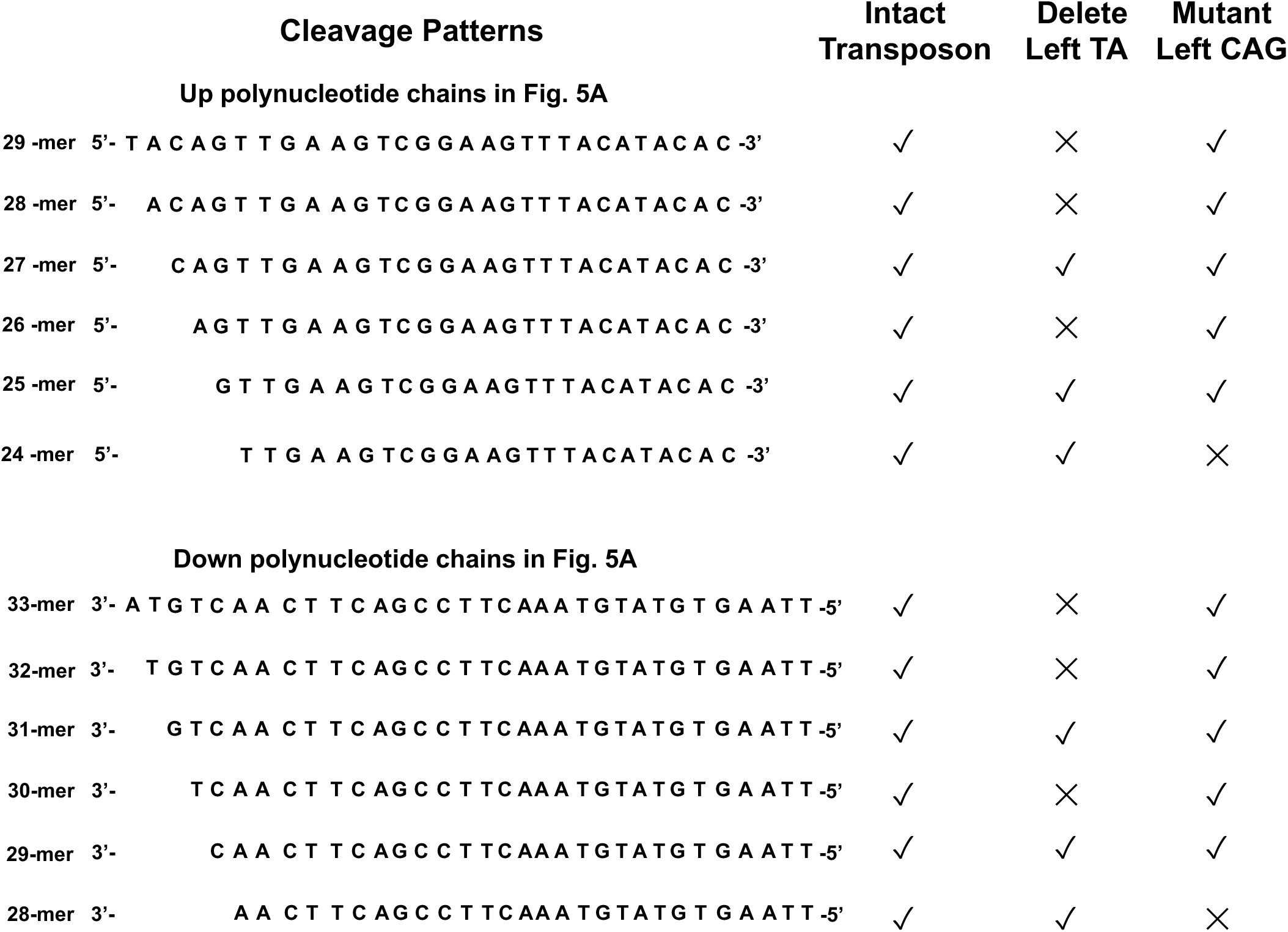
Cleavage patterns within the left end of transposons.

**Figure S5.**
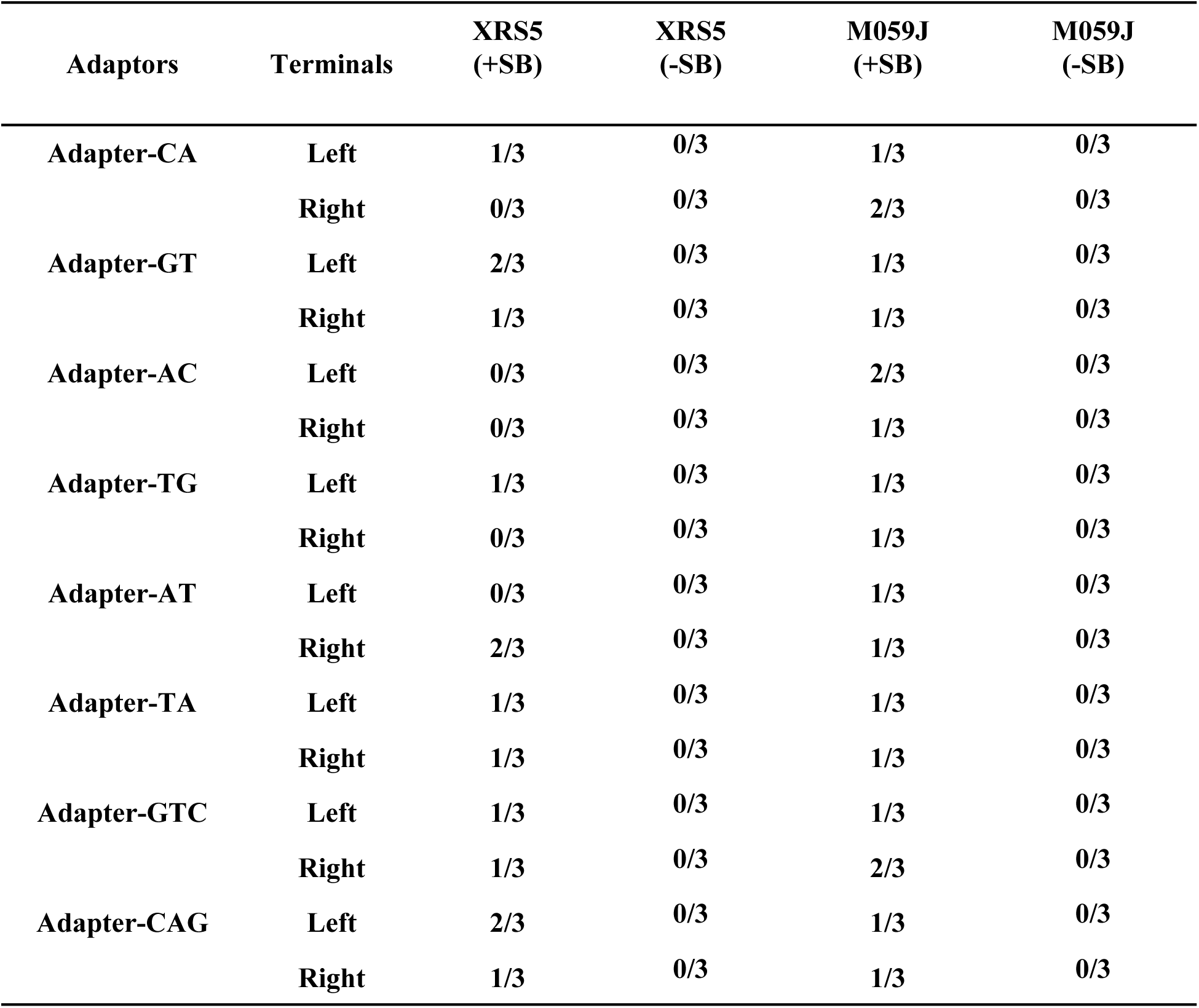
Terminal structures in NHEJ deficiency cells. XRS5 and M059J cells were transfected with pT/SVneo ± pCMV-SB11 (+SB and –SB). Total DNA was extracted and ligated to appropriate adaptors. Adaptor-mediated PCR was performed to generate products from the left or right termini for sequencing. PCR^+^/Exps indicates the frequency of positive PCR products (PCR^+^) containing the precise terminal sequence from six independent experiments (Exps).

**Figure S6.**
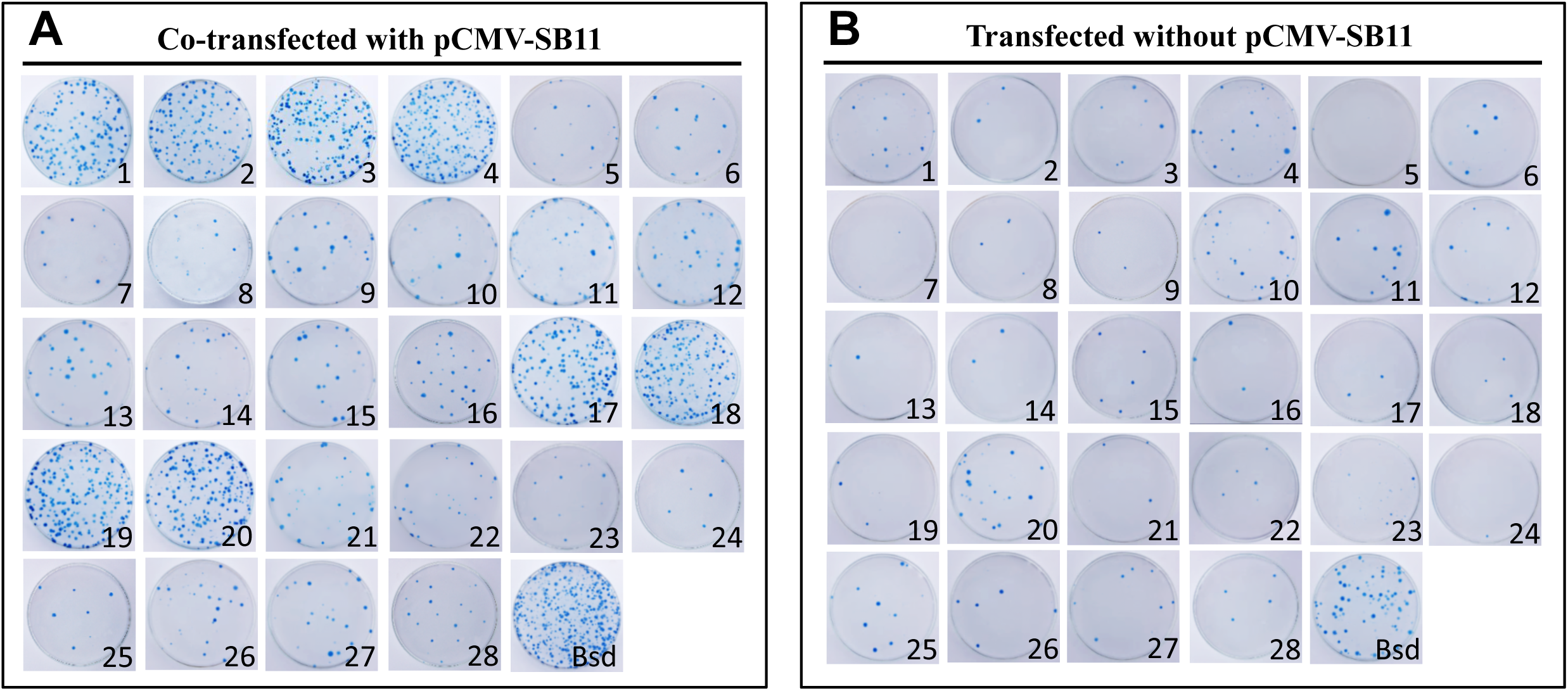
Transposition rates of artificial SB ETFs. Artificial SB ETFs were co-transfected with (**A**) or without (**B**) pCMV-SB11 into HeLa cells. The Bsd-resistant cell colonies were fixed with formaldehyde, stained with methylene blue and counted, and photographed. Unmodified plasmid pT_2_/zbHsp70-Bsd-SV40 (marked as Bsd) was used as the positive control.

**Table S1.**
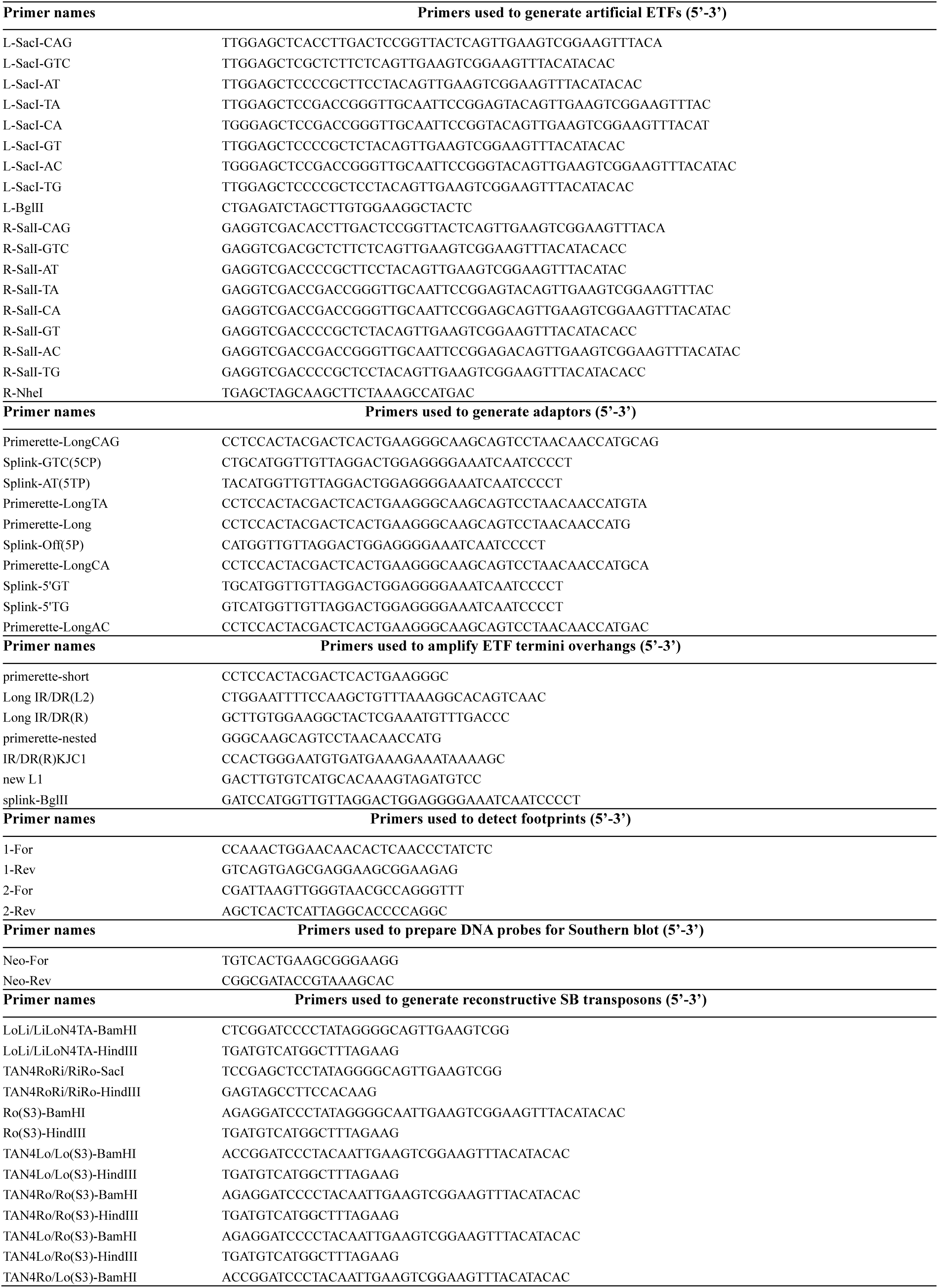
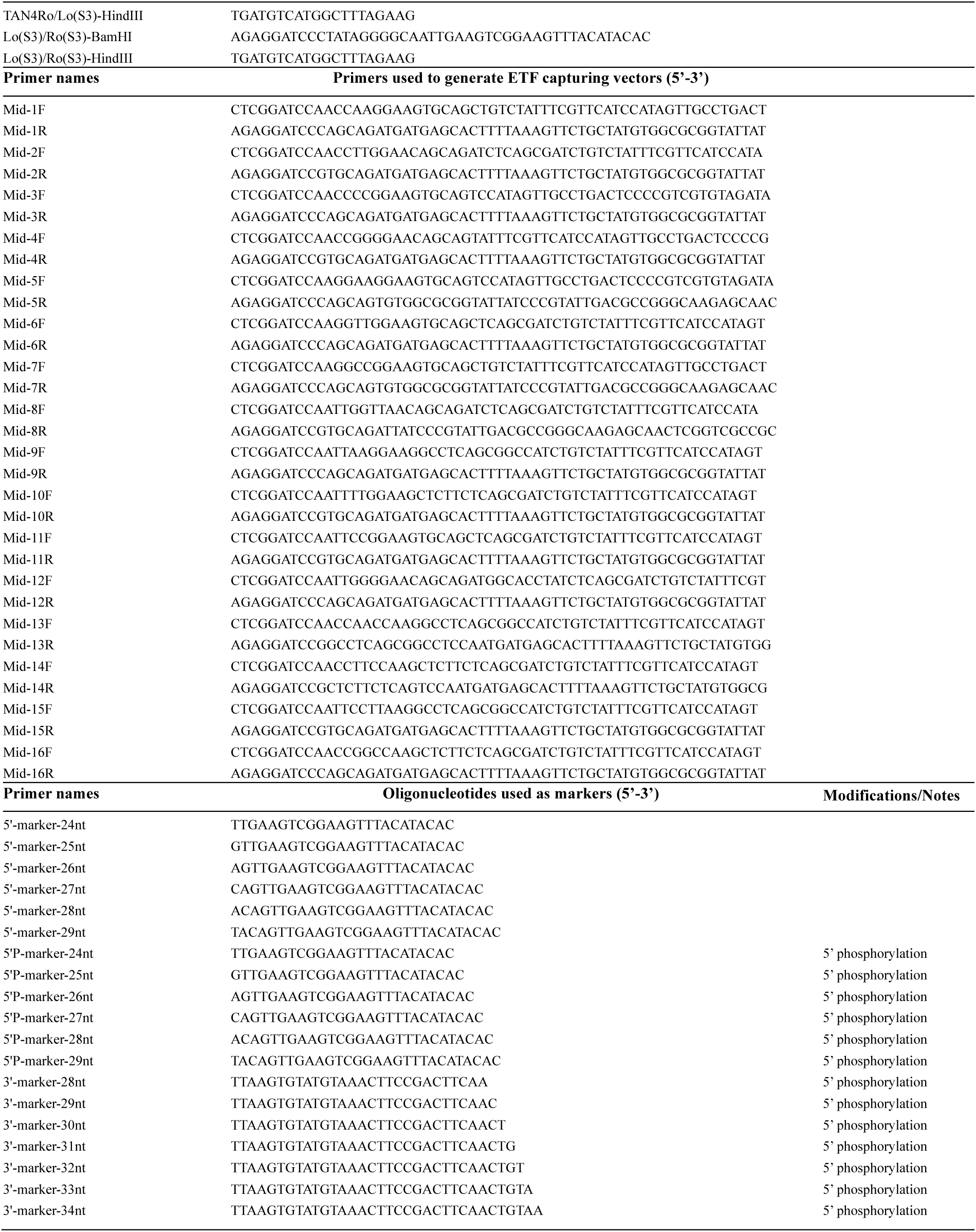
Oligos used in this study

**Table S2.**
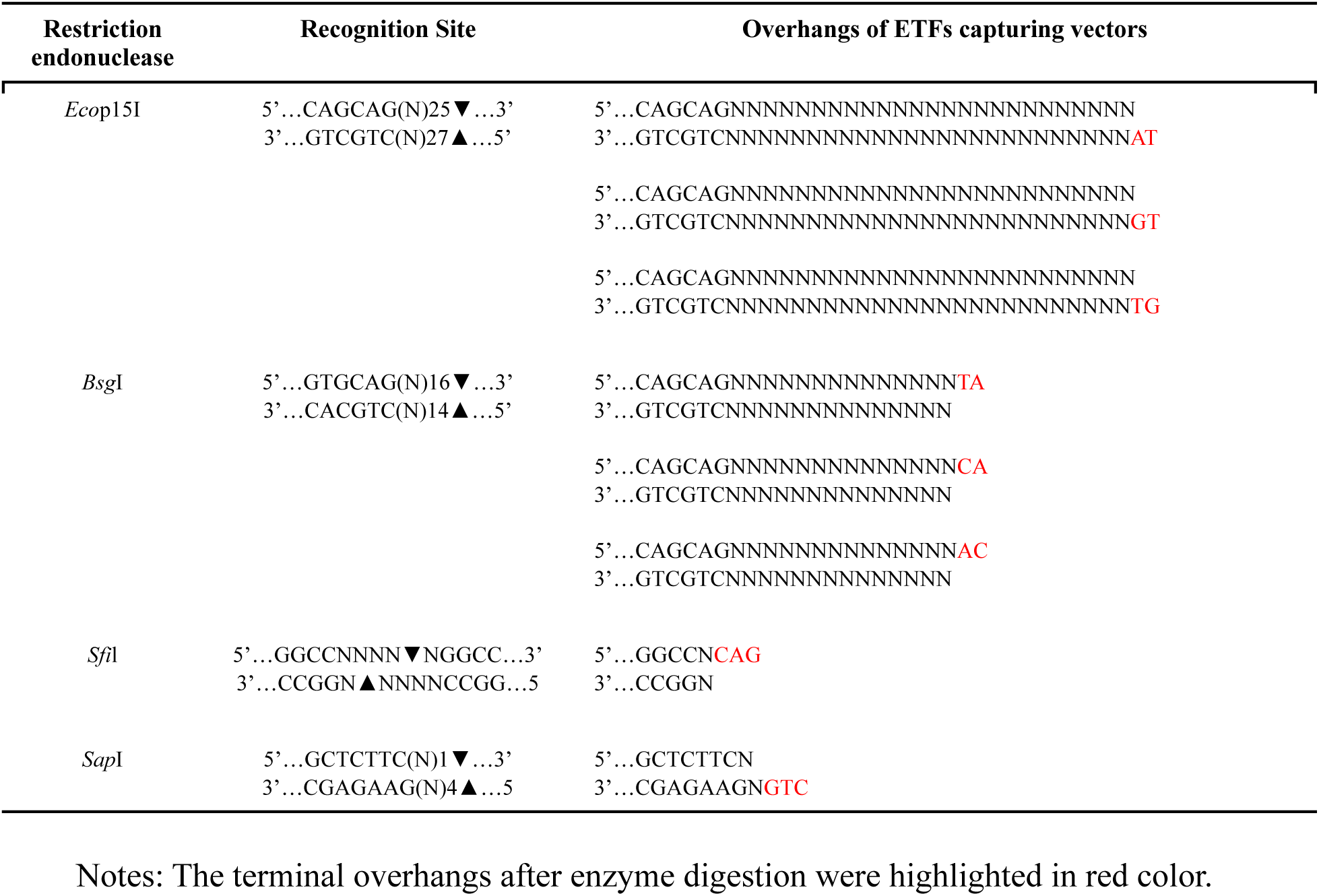
Restriction endonucleases used to generate the terminal overhangs of ETFs capturing vectors

**Table S3.**
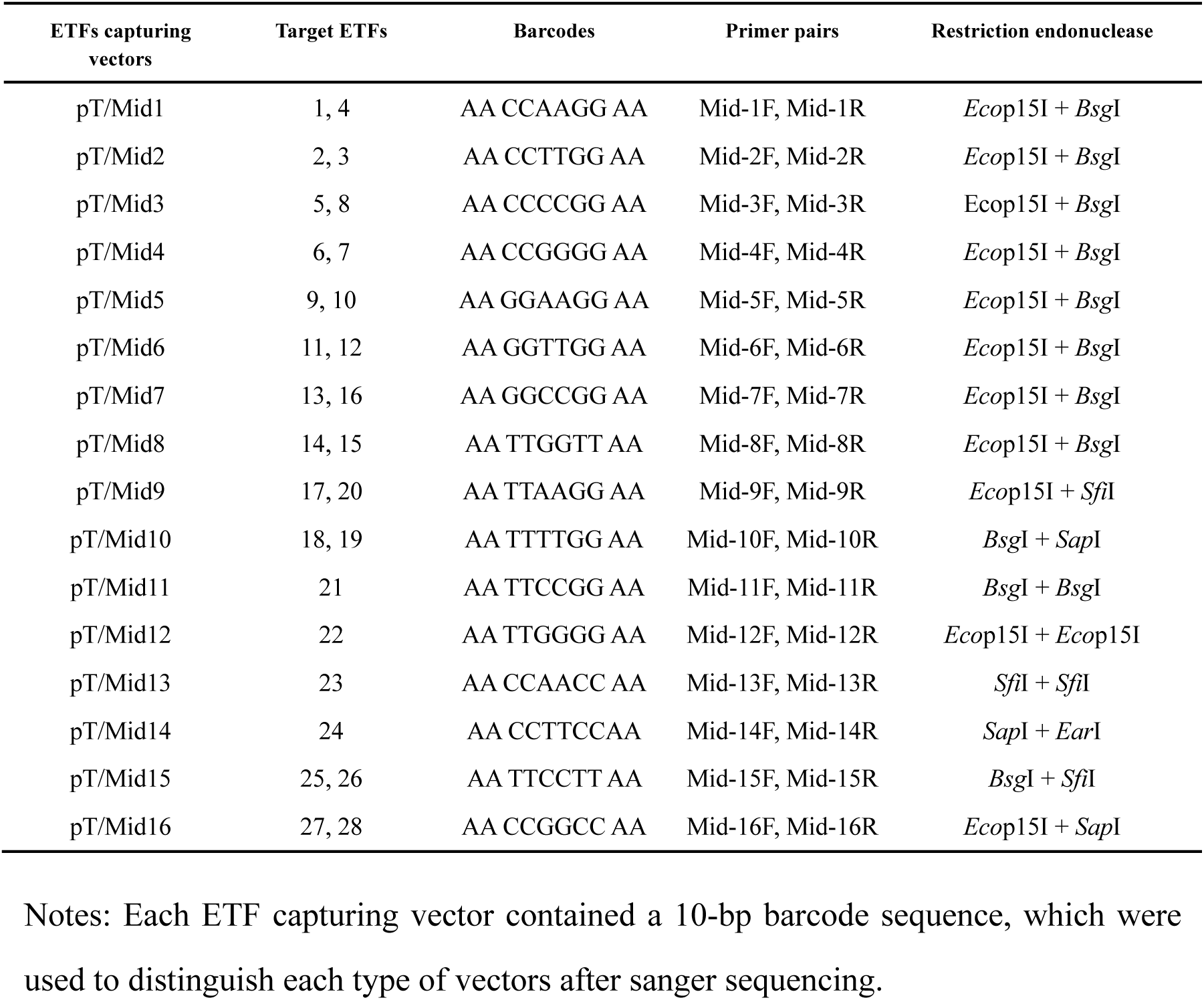
Generation of the ETFs capturing vectors

**Table S4.** Restriction endonucleases used to generate artificial ETFs

| Restriction endonuclease | Recognition Site | Overhangs of artificial ETFs |
| --- | --- | --- |
| *FauI* | 5'...CCCGC(N)4▼...3'<br>3'...GGGCG(N)6▲...5' | 5'...CCCGCNNNN<br>3'...GGGCGNNNN**AT**<br>5'...CCCGCNNNN<br>3'...GGGCGNNNN**GT**<br>5'...CCCGCNNNN<br>3'...GGGCGNNNN**TG** |
| *BcgI* | 5'...▼10(N)CGA(N)6TGC(N)12▼...3'<br>3'...▲12(N)GCT(N)6ACG(N)10▲...5' | 5'...10(N)CGA(N)6TGCNNNNNNNNNN**TA**<br>3'...12(N)GCT(N)6ACGNNNNNNNNNN<br>5'...10(N)CGA(N)6TGCNNNNNNNNNN**CA**<br>3'...12(N)GCT(N)6ACGNNNNNNNNNN<br>5'...10(N)CGA(N)6TGCNNNNNNNNNN**AC**<br>3'...12(N)GCT(N)6ACGNNNNNNNNNN |
| *BsaXI* | 5'...▼9(N)AC(N)5CTCC(N)10▼...3'<br>3'...▲12(N)TG(N)5GAGG(N)7▲...5' | 5'...9(N)AC(N)5CTCCNNNNNNNN**CAG**<br>3'...12(N)TG(N)5GAGGNNNNNNNN |
| *BspQI* | 5'...GCTCTTC(N)1▼...3'<br>3'...CGAGAAG(N)4▲...5' | 5'...GCTCTTCN<br>3'...CGAGAAGN**GTC** |

Notes: The terminal overhangs after enzyme digestion were highlighted in red color.

**Table S5.**
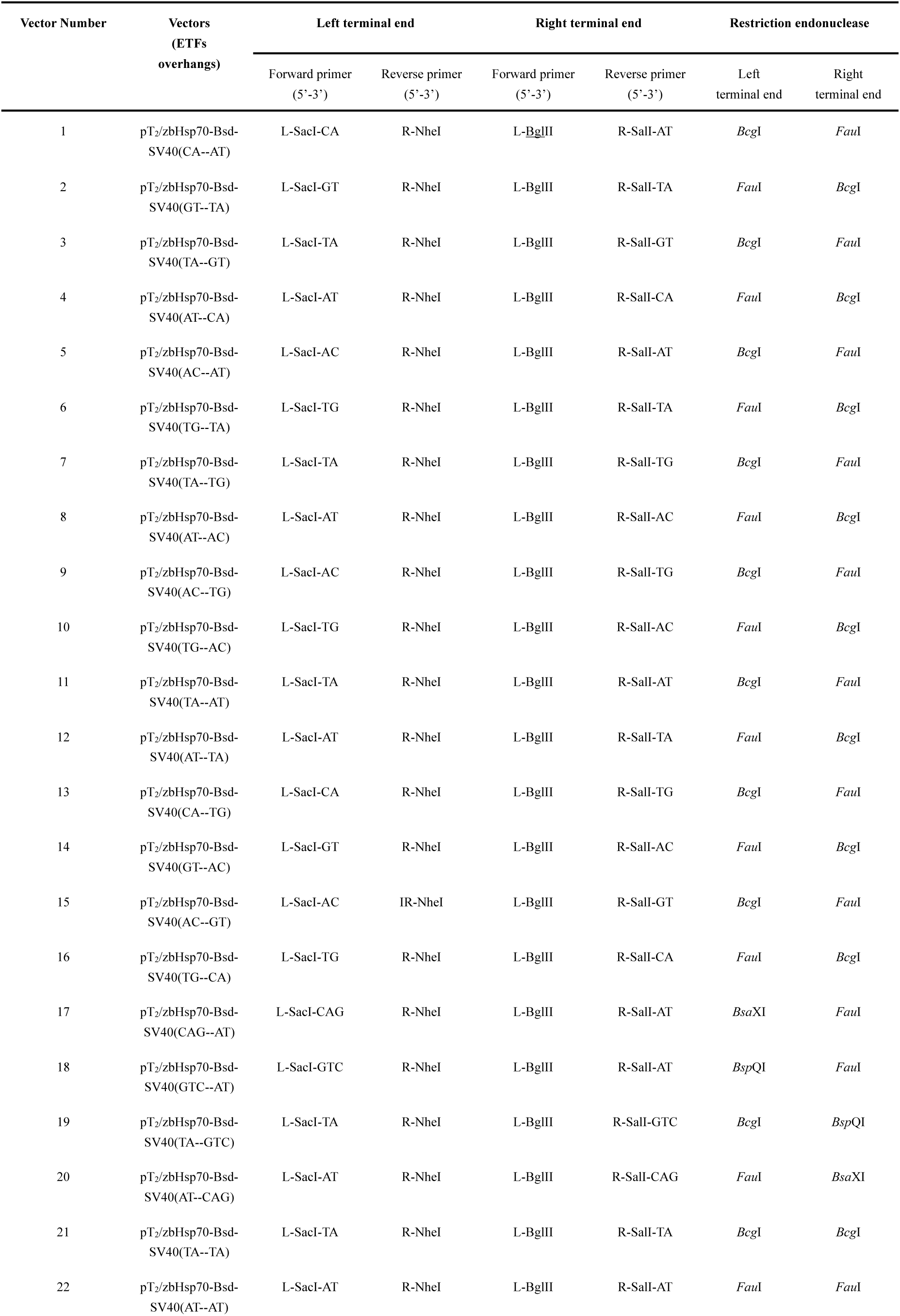
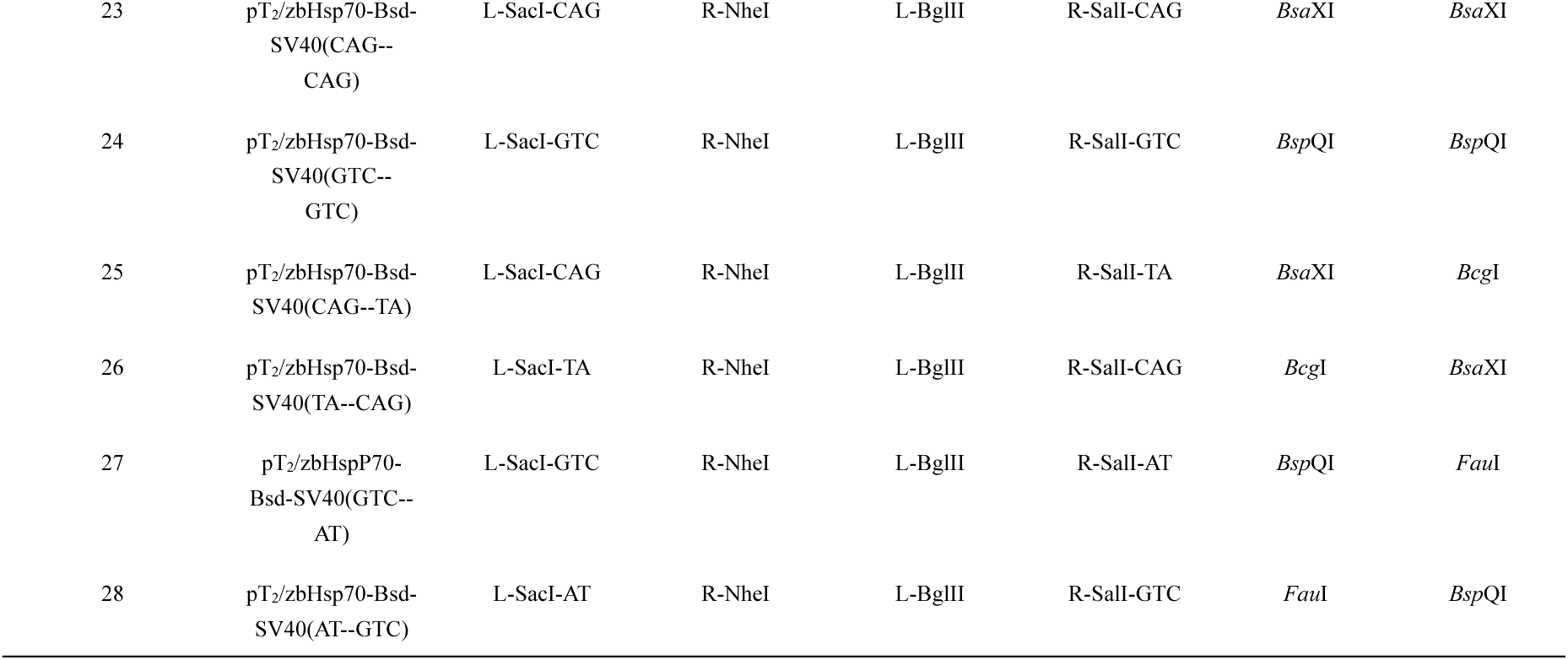
Primer pairs used to construct vectors containing artificial ETFs

**Table S6.** Oligos used for formation of adapters

| **Adapters** | **Forward oligo (Primerettes)** | **Reverse oligo (Splinks)** |
| --- | --- | --- |
| Adapter-CA | Primerette-LongCA | Splink-Off(5P) |
| Adapter-GT | Primerette-Long | Splink-5'GT |
| Adapter-AC | Primerette-LongAC | Splink-Off(5P) |
| Adapter-TG | Primerette-Long | Splink-5'TG |
| Adapter-AT | Primerette-Long | Splink-AT(5TP) |
| Adapter-TA | Primerette-LongTA | Splink-Off(5P) |
| Adapter-GTC | Primerette-Long | Splink-GTC(5CP) |
| Adapter-CAG | Primerette-LongCAG | Splink-Off(5P) |
| Adapter-BglII | Primerette-Long | Splink-BglII |

